# Seizures are a druggable mechanistic link between TBI and subsequent tauopathy

**DOI:** 10.1101/2020.05.12.091819

**Authors:** Hadeel Alyenbaawi, Richard Kanyo, Laszlo F. Locskai, Razieh Kamali-Jamil, Michèle G. DuVal, Qing Bai, Holger Wille, Edward A. Burton, W. Ted Allison

## Abstract

Traumatic brain injury (TBI) is a prominent risk factor for neurodegenerative diseases and dementias including chronic traumatic encephalopathy (CTE). TBI and CTE, like all tauopathies, are characterized by accumulation of Tau into aggregates that progressively spread to other brain regions in a prion-like manner. The mechanisms that promote spreading and cellular uptake of tau seeds after TBI are not fully understood, in part due to lack of tractable animal models. Here, we test the putative roles for excess neuronal activity and dynamin-dependent endocytosis in promoting the *in vivo* spread of tauopathy. We introduce ‘tauopathy reporter’ zebrafish expressing a genetically-encoded fluorescent Tau biosensor that reliably reports accumulation of human tau species when seeded *via* intra-ventricular brain injections. Subjecting zebrafish larvae to a novel TBI paradigm produced various TBI symptoms including cell death, hemorrhage, blood flow abnormalities, post–traumatic seizures, and Tau inclusions. Bath application of anticonvulsant drugs rescued TBI-induced tauopathy and cell death; these benefits were attributable to inhibition of post-traumatic seizures because co-application of convulsants reversed these beneficial effects. However, one convulsant drug, 4-Aminopyridine, unexpectedly abrogated TBI-induced tauopathy - this was due to its inhibitory action on endocytosis as confirmed via additional dynamin inhibitors. These data suggest a role for seizure activity and dynamin-dependent endocytosis in the prion-like seeding and spreading of tauopathy following TBI. Further work is warranted regarding anti-convulsants that dampen post-traumatic seizures as a route to moderating subsequent tauopathy. Moreover, the data highlight the utility of deploying *in vivo* Tau biosensor and TBI methods in larval zebrafish, especially regarding drug screening and intervention.

**Graphical Abstract:** 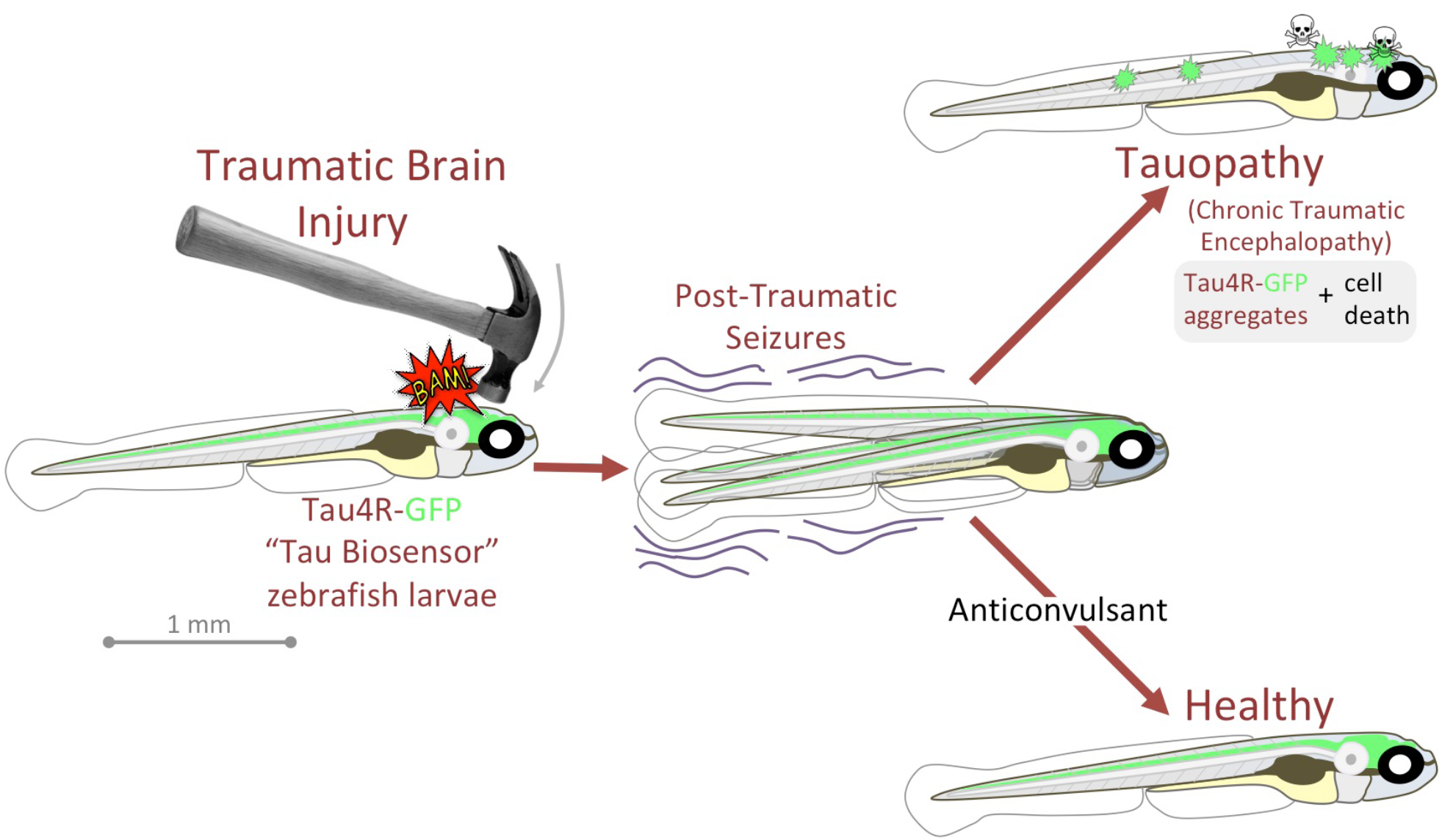

**Highlights:**

- Introduces first Traumatic Brain Injury (TBI) model in larval zebrafish, and its easy
- TBI induces clinically relevant cell death, haemorrhage & post-traumatic seizures
- Ca^2+^ imaging *during* TBI reveals spike in brain activity concomitant with seizures
- Tau-GFP Biosensor allows repeated *in vivo* measures of prion-like tau aggregation
- post-TBI, anticonvulsants stop tauopathies akin to Chronic Traumatic Encephalopathy

## Introduction

Traumatic brain injury (TBI) is a leading cause of mortality and disability worldwide (Hay et al., 2016; Nguyen et al., 2016; Rimel et al., 1981). It also is a prominent risk factor for neurodegeneration and dementia, such as chronic traumatic encephalopathy (CTE) (Chauhan, 2014; Gardner and Yaffe, 2015; Uryu et al., 2007). TBI can result from direct physical insults, from rapid acceleration and deceleration of the brain, or from shock wave impacts such as explosive blasts (Cruz-Haces et al., 2017). Regardless, the primary mechanisms have much in common and the neuropathology in TBI and CTE patients includes the wide distribution of hyperphosphorylated tau pathology, axonal degeneration, and neuronal loss (Hay et al., 2016; Johnson et al., 2013; McKee et al., 2015; Ojo et al., 2016). The mechanisms whereby physical injury is translated into progressive tau pathology remain unresolved and represent prospective therapeutic targets. Progress on this front is hampered by lack of access to suitable models: applying physical injury to a cell culture is difficult and poorly represents the complex biopathology that intertwines many multifaceted aspects of brain physiology.

The progressive deposition of hyperphosphorylated tau protein in filamentous forms is a defining hallmark of tauopathies, which includes Alzheimer’s disease (AD), CTE, and several other dementias. Each of the tauopathies affects distinct brain regions and has a unique clinical presentation (Kovacs, 2017; Orr et al., 2017). Early in CTE, hyperphosphorylated tau is accumulated in a cluster of perivascular neurons and glia in the depths of cortical sulci. Later in CTE, tau pathology is widespread and incorporates cortical and subcortical grey-matter areas (Hay et al., 2016; Johnson et al., 2012; McKee et al., 2015). This broad spreading of tau pathology in CTE can also be observed following TBI ascribed to single trauma events (Johnson et al., 2012). This spreading of tauopathy is consistent with a prion-like mechanism; indeed brain homogenates from mice subjected to TBI can initiate p-Tau pathology when injected into healthy wildtype mice (Zanier et al., 2018). The recipient mice develop a p-tau pathology similar to single severe TBI patients, which then spreads from injection sites to distant regions, behaving similarly to bona fide prions (Zanier et al., 2018).

Beyond TBI, the self-propagation and prion-like spread of tau aggregates is thought to play a key role in the progression of other tauopathies such as AD (Iba et al., 2013; Iba et al., 2015; Mudher et al., 2017; Narasimhan et al., 2017; Sanders et al., 2014). Mechanisms of tau spreading, and the therapeutic targets they offer, have principally been defined *in vitro* and include tunnelling nanotubes and extracellular vesicles (EVs such as exosomes and synaptic vesicles) and their uptake via endocytosis (Colin et al., 2020; Demaegd et al., 2018; Evans et al., 2018). In AD and other tauopathies, observations from patients and mice have highlighted the capacity of tau seeds to spread trans-synaptically (Goedert et al., 1989; Pickett et al., 2017). Moreover, it has been shown that neuronal activity serves an important role in the spread of tau pathology and general proteostasis (Pickett et al., 2017; Wu et al., 2016; Yamada et al., 2014). Stimulation of neuronal activity increased the extracellular release of tau to the media *in vitro*, and enhanced tau pathology in a mouse model of familial frontotemporal dementia (Pickett et al., 2017; Wu et al., 2016; Yamada et al., 2014). Whether similar mechanisms of tau release and spread occur following TBI remains unknown.

In this light, an intriguing aspect of TBI is the prominence of post-traumatic seizures that might be predicted to initiate the aggregation and/or exacerbate the spread of tau pathology. Seizures are one of the key consequences of all types of TBI, and they have been more commonly reported in patients who suffered from blast injuries (Asikainen et al., 1999; Salinsky et al., 2015). Though the exact prevalence remains undetermined (Lucke-Wold et al., 2015), it is anticipated that over 50% of TBI patients with severe injuries develop seizures or post-traumatic epilepsy (Kovacs et al., 2014). A link between seizures and tau pathology is suggested by increased prevalence of seizures in AD patients and animal models of AD (Sanchez et al., 2018; Yan et al., 2012). Whether reducing post-traumatic seizures can delay or minimize the progression of tauopathy has yet to be fully explored.

This knowledge gap is due in part to a lack of accessible *in vivo* models that can report the progression and spread of tauopathy, or that allow neural activity associated with TBI to be measured and manipulated. To address these issues, we engineered a tauopathy biosensor transgenic zebrafish that develops GFP+ puncta when tau aggregates within the brain or spinal cord. Additionally, we introduce a simple medium-throughput method to induce TBI in zebrafish larvae. Combining these novel approaches, we found that post-traumatic seizures correlate strongly with spreading tau pathology following TBI. Manipulating this seizure activity mitigated tau aggregation and revealed a critical role for endocytosis in the prion-like spread of tau seeds *in vivo* following TBI. The results from our novel *in vivo* TBI model implicate seizures and dynamin-dependent endocytosis in the spread of tau seeds, thereby offering potential therapeutic targets.

## Methods

### Animal Ethics and Zebrafish Husbandry

Zebrafish were raised and maintained following protocol AUP00000077 approved by the Animal Care and Use Committee: Biosciences at the University of Alberta, operating under the guidelines of the Canadian Council of Animal Care. The fish were raised and maintained within the University of Alberta fish facility under a 14/10 light/dark cycle at 28°C as previously described (Westerfield, 2000).

### Generating Transgenic Tauopathy Reporter Zebrafish

To engineer the transgenic Tau4R-GFP reporter zebrafish, the human wild-type MAPT sequence of the four-microtubule binding repeat domain (aa 244-372 of the full-length TAU 2N4R, NCBI NC_000017.11, protein id NP_005901) with a seven-amino acid C-terminal linker (RSIAGPA) was ordered as a gene block from IDT. The gene block was subcloned into a middle entry cloning vector (Multisite Gateway® technology, ThermoFisher). This was recombined with the p5E-enolase2 and p3E-GFP components into destination vector pDestTol2CG2 of the Tol2kit (Guo and Lee, 2011; Kwan et al., 2007). The destination vector contained a reporter construct [encompassing EGFP driven by the cardiac myosin light chain (clmc) promoter that helps identify stable transgenic zebrafish]. The resulting plasmid pDestTol2CG2.eno2:Tau4R-GFP was delivered in a 10μl injection solution including 750ng/μl of the construct mixed with 250ng/μl Tol2 transposase mRNA, 1μl of 0.1M KCL, and 20% phenol red. The solution was injected into the single-cell embryos of Casper zebrafish line (transparent zebrafish line)(White et al., 2008). Injected embryos were screened for mosaic expression of the Tau4R-GFP transgene at two days post-fertilization (dpf) using a Leica M165 FC dissecting microscope. F0 mosaic fish were raised to adulthood and outcrossed. Successful F1 embryos were identified by their abundant expression of Tau4R-GFP in the CNS and the green heart marker. The stable transgenic line *Tg(eno2:Hsa.MAPT_Q244-E372-EGFP)*^*ua3171*^ was assigned the allele number ua3171.

An equivalent transgenic zebrafish biosensor was engineered to detect human SOD1 aggregation. Subcloning from existing vectors (Pokrishevsky et al., 2018) produced pDestTol2CG2.eno2:SOD1-GFP and similar transgenesis methods engineered the *Tg[eno2:SOD1-GFP]* zebrafish line that was assigned allele number ua3181.

### Cell culture and Generation of Tauopathy Reporter Stable Cell Line

To move the Tau4R-GFP reporter above into a vector appropriate for cell culture, BamHI and Xhol nuclease restriction enzymes were employed to remove the Tau4R-GFP fragment from pDest tol2CG2.eno2.Tau4R-GFP.pA. The Tau4R-GFP fragment was subcloned into the pCDNA3.1 vector using a T4 DNA ligase enzyme. Sequencing of the cloned vector with the following reverse primer for GFP (TCTCGTTGGGGTCTTTGCTC) confirmed the proper orientation. Purification of the plasmid was conducted with the Qiagen purification kit. HEK293T cells were grown in Dulbecco’s modified Eagle’s medium (GibcoTM, ThermoFisher) supplemented with 10% fetal bovine serum and 1% penicillin/streptomycin. All cells were maintained at 37°C in a humidified 5% CO2 incubator. For passaging cells, cells were washed with phosphate-buffered saline (PBS) before trypsinization with 0.05 Trypsin-EDTA (Sigma Aldrich, T4174).

HEK293T cells were plated at 1 × 10^6^ cells/well in six-well plates. Cells were transfected with pcDNA3.1.Tau4R-GFP plasmid 24 hours after plating using lipofectamine 2000 reagents according to the manufacturer’s guidelines. Briefly, 4µg of pcDNA3.1.Tau4R-GFP was diluted in 250µl of Opti-MEM media (GibcoTM, ThermoFisher). The expression of the fluorescent reporter was confirmed the next day through microscopic analysis. A stable cell line was established by replating the transfected cells at a 1:10 dilution and selecting in DMEM media containing 1200μg/ml geneticin (GibcoTM, ThermoFisher). Expression of the fused fluorescent proteins in the stable cell lines was confirmed using fluorescent microscopy. Polyclonal cells and monoclonal cells were grown to confluency in 10cm dishes, then stored in liquid nitrogen until use.

### Immunoblotting of Cell Lysate and Zebrafish Brain Lysate

For cell lysate preparation, cells were washed with cold PBS, then collected and incubated with cold lysis buffer (150mM NaCl, 50mM Tris-HCl (pH 8), 1 mM EDTA and 1% Nonidet P-40) supplemented with protease inhibitor (Cocktail Set III; Millipore) for 10 mins on ice. Cells were lysed using a bio-vortexer homogenizer for 20 sec for two rounds. The lysate was centrifuged at 13000rpm for 10 mins at 4°C. The supernatant was collected, and the protein concentration was determined using the Qubit™ Protein Assay Kit (Invitrogen).

For zebrafish brain lysate preparation, the brains of adult zebrafish were dissected. Brains were homogenized in cell lysis buffer (20mM HEPES, 0.2mM EDTA, 10mM NaCl, 1.5mM MgCl2, 20% glycerol, 0.1% Triton-X) with protease inhibitor and phospSTOP (Sigma-Aldrich) in the case of pt406 Tg. Brains were lysed using a bio-vortexer homogenizer and sonicated for 3 sec for one round. Samples were centrifuged as above and concentration of the samples was assessed in a Qubit® fluorometer (Invitrogen).

For immunoblotting, 30-40µg of the total protein was combined with 2X sample buffer (Sigma-Aldrich) and boiled for 10 mins before loading in 11% SDS-PAGE. Electrophoresis was performed using the Bio-Rad Power PAC system in running buffer (25 mM Tris base, 192 mM glycine and 0.1% SDS). The gel was transferred to a PVDF membrane using a wet transfer system. All membranes were blocked for one hour in protein-free blocking buffer PBS (ThermoFisher) or TBST with 5% milk and then incubated with primary antibody overnight at 4°C with gentle agitation. The primary antibodies used in this study include rabbit monoclonal GFP (abcam, EPR14104) at 1:3000 dilution, rabbit anti-β-actin (Sigma-Aldrich, A2066) at 1:10000. All membranes were washed three times with 1X TBST before incubation with secondary antibody (goat-anti-mouse HRP or HRP-conjugated anti-rabbit at 1:5000 dilution (Jackson ImmunoResearch) for one hour at room temperature. The membranes were washed for the final time before visualization using Pierce® ECL Western Blotting Substrate (ThermoFisher) on a ChemiDoc (Biorad). For stripping and re-probing, the membranes were stripped using mild stripping buffer (199.8 mM Glycine, 0.1% SDS and 1% Tween 20 with a pH of 2.2) before blocking them and repeating the methods described before.

### Immunohistochemistry

Larvae were fixed overnight in 4% paraformaldehyde, either one day after being subjected to TBI or following the subsequent application of drugs as indicated. Immunostaining of Activated-Caspase3 on whole-mount larvae was carried out as previously described (DuVal et al., 2014). Larvae were washed with 0.1 M PO4 with 5% sucrose three times before washing with 1% Tween in H_2_O (pH 7.4), and then -20°C acetone. Larvae were incubated in PBS3+ containing 10% normal goat serum for one hour and then incubated with primary antibody with 2% normal goat serum in PBS3+. The primary antibody used was polyclonal Anti-Active-Caspase-3 (BD Pharmingen, 559565) at 1:500 dilution. The secondary antibody applied was Alexafluor 647 anti-rabbit at 1:200 dilution (Invitrogen). Larvae were counterstained with 4’,6-diamidino-2-phenylindole (DAPI) (ThermoFisher) for 30 minutes.

### Preparations of Mouse Brain Homogenate (Crude and PTA Precipitated)

Brains from TgTau^P301L^ mice and non-Tg littermate controls (129/SvEvTac genetic background) were provided by Dr. David Westaway and Dr. Nathalie Daude (Eskandari-Sedighi et al., 2017; Murakami et al., 2006). Crude brain homogenate was prepared by homogenizing the brains to 10% (wt/vol) in calcium- and magnesium-free DPBS that included a protease inhibitor and phosSTOP, using a glass homogenizer and power gen homogenizer (Fisher Scientific). Samples were then centrifuged at 13000 rpm for 15 mins at 4°C. The clear supernatant was collected, aliquoted and stored in -80°C until use for experiments.

The phosphotungstate anion (PTA)-precipitated brain homogenate was prepared as described (Woerman et al., 2016). Briefly, 10% (wt/vol) brain homogenate was prepared as reported above and mixed with a final concentration of 2% sarkosyl (Sigma Aldrich) and 0.5% benzonase (Sigma Aldrich, E1014), and then incubated at 37°C for two hours with constant agitation in an orbital shaker. Sodium PTA (Sigma Aldrich) was dissolved in ddH_2_O, and the pH was adjusted to 7.0 before it was added to the samples at a final concentration of 2% (vol/vol). The samples were then incubated overnight under the previous conditions. The next day, the samples were centrifuged at 16,000g for 30 mins at room temperature. The supernatant was discarded, while the resulting pellet was resuspended in 2% (vol/vol) PTA in ddH_2_O (pH 7.0) and 2% sarkosyl in DPBS. The samples were next incubated for one hour before the second centrifugation. The supernatant was removed and the pellet was re-suspended in DPBS. An aliquot of 5μl of PTA purified brain homogenate was employed for electron microscopy (EM) analysis to confirm the presence of fibrils in each sample.

### Tau Fibrillization and EM Analysis

Synthetic human tau protein (wildtype full-length monomers) was purchased as a lyophilized powder (rPeptide, T-1001-2) and resuspended in ddH_2_O at a concentration of 2mg/ml. The recombinant protein was fibrillized as described previously (Guo and Lee, 2011). Recombinant tau was incubated with 40μM low-molecular-weight heparin and 2mM DTT in 100 mM sodium acetate buffer (pH 7.0) at 37°C, thereafter being agitated for seven days. The fibrillization mixture was centrifuged at 50,000g for 30 mins, and the resulted pellet was resuspended in 100 mM sodium acetate buffer (pH 7.0) without heparin or DTT. Successful fibrillization was verified by EM.

Negative staining for EM analysis of fibrils was conducted as described elsewhere (Eskandari-Sedighi et al., 2017). Briefly, 400 mesh carbon-coated copper grids (Electron Microscopy Sciences) were glow-discharged for 40 sec before adding the sample aliquots. PTA-purified brain homogenates or synthetic tau fibrils (5µL) were applied on the top of the grid for 1 min. These grids were washed using 50µL each of 0.1M and 0.01M ammonium acetate and negatively stained with 2 × 50µL of filtered 2% uranyl acetate. After removing excess stain and drying, the grids were examined with a Tecnai G20 transmission electron microscope (FEI Company) with an acceleration voltage of 200 kV. Electron micrographs were recorded with an Eagle 4k × 4k CCD camera (FEI Company).

### Liposome-Mediated Transduction of Brain Homogenate into Tauopathy Reporter Cells

Polyclonal Tau4R-GFP cells were plated at 2 × 10^5^ per well in 24-well plates. Cells were transduced the next day, using 40µl of 10% clarified brain homogenate combined with Opti-MEM to a final volume of 50µl. A further 48µl of Opti-MEM and 2µl of Lipofectamine-2000 (Invitrogen) was added to the previous Opti-MEM mixture to a total volume of 100µl and incubated for 20 mins. The liposome mixture was applied to the cells for 18 hours, and cells were then washed with PBS, trypsinized, and re-plated on coated coverslips (ThermoFisher) for imaging and analysis.

For PTA-precipitated brain homogenate, 1:10 dilution of precipitated fibrils was used for the transfection. 5µl of PTA-purified fibrils was diluted in 45µl Opti-MEM to a final volume of 50µl. The previous Opti-MEM mixture was added to 47µl of Opti-MEM and 3µl of Lipofectamine-2000 and incubated in room temperature for 2 hours as described in (Safar et al., 1998; Woerman et al., 2016). The mixture was added to cells, washed after 18 hours and re-plated before analysis exactly as mentioned previously.

### Quantification of the Percentage of Cells with Positive Inclusion

Prior to imaging, transfected cells were fixed 2% PFA in PBS for 15 mins. Samples were then washed twice with PBS then stained with DAPI (1:3000 from 1mg/ml stock) for six mins. Cells were imaged using a Zeiss LSM 700 scanning confocal microscope featuring Zen 2010 software (Carl Zeiss, Oberkochen, Germany). Due to increased brightness of the GFP+ puncta formed after introduction of brain homogenate, GFP exposure was minimized for those cells only. To quantify the GFP+ puncta, a total of nine images were collected and analyzed for each condition, each with ∼100 cells. DAPI-positive nuclei were utilized to determine the number of cells per image. The number of cells with inclusions (multiple nuclear inclusions or one cytoplasmic puncta) were counted and the percentage was calculated.

### Brain Ventricle Injections into Tauopathy Reporter Larvae

Injections into the larval zebrafish brain (intraventricular space) were performed as described previously with few modifications (Gutzman and Sive, 2009). Embryos at 2 dpf (days post-fertilization) were removed from their chorions and anesthetized with 4% tricaine (MS-222, Sigma Aldrich). The embryos were placed in a 1% agarose-coated dish with small holes. Under a stereomicroscope, the immobilized embryos were oriented so that the brain ventricles were accessible for injections. The injection was carried out via pulled capillary tubes mounted in a micromanipulator. The injection volume was calibrated to 5nL by injection into mineral oil and measurement with an ocular micrometer. Thereafter, the needle containing the injection solutions was placed through the roof plate of the hindbrain and 5-10nL of either 10% clarified brain homogenate (TgTau^P301L^ mice or wildtype littermate control), or synthetic tau, were mixed with 20% dextran Texas Red fluorescent dye (Invitrogen) and injected into the ventricles. For all the brain injection experiments, an uninjected control group and control group injected only with 20% red dextran fluorescent dye in PBS were included. After the injections, embryos were screened using a Leica M165 FC dissecting microscope and appropriately injected larvae were gathered for further analysis. The injections were considered appropriate if they had sharp edges and non-diffuse dye in the ventricle (Fig. S2A). Larvae receiving improper injections, in which the needle was inserted too deep in the brain ventricles resulting in the dye being visible outside the ventricle space and/or in the yolk, were excluded from analysis.

### Microscopy Analysis of GFP Positive Puncta in Tau Reporter Larvae

For the microscopic analysis of GFP-positive inclusions, larvae that were either injected or treated with traumatic injury, along with the control groups, were anesthetized via tricaine at the indicated time point (two, three, four, or five days post-injection (dpi) or post traumatic injury (dpti) depending on the experiment). Images for GFP-positive puncta on the brain area or lateral line above the spinal cord were taken using a Leica M165 FC dissecting microscope and the number of GFP-positive puncta were manually counted.

### Traumatic Brain Injury (TBI) paradigm for Zebrafish Larvae

To induce TBI, 10-12 unanesthetized larvae (3 dpf) were loaded into a 10ml syringe with 1ml of E3 media. The syringe was blocked using a stopper valve to ensure no larvae or media left the syringe upon compression of the plunger. The syringe was held vertical using a metal tube holder at the bottom end of a 48” tube apparatus. A defined weight (between 30 and 300g) was dropped manually from the top of the tube. The tube diameter was matched to (slightly greater than) the weight’s diameter to enhance repeatability. This was either done once or repeated three times, with either 65 or 300g weights. Once larvae were subjected to the traumatic brain injury, they were moved back to a petri dish with fresh media and maintained for further analysis.

### Quantifying the pressure induced during TBI

To characterize the dynamic changes in pressure that occurred within the syringe during the TBI events, the stopper valve attached to the syringe (described immediately above) was replaced with a piezoresistive pressure transducer (#MLT844 AD Instruments, Colorado Springs, CO). Events were monitored via a PowerLab 2/26 data acquisition device and LabChart 7 software (AD Instruments). The pressure transducer was zeroed to report gauge pressure (pressure changes relative to atmospheric pressure) and was calibrated against a manometer (Fisherbrand Traceable from Thermoscientific, Ottawa ON). After each weight drop, the syringe apparatus was reset to remove any air bubbles and the pressure transducer was zeroed. Time courses of induced pressure were reported over a 350 msec time frame with 50 msec of base line recording, while mean and maximum pressure values were calculated from the initial 300 msec following the impact of the weight.

### Recording blood flow following TBI

Abnormalities of blood flow and circulation resulted from TBI was detected 5 to 10 mins after larvae was subjected to TBI. The blood flow in the tail area of zebrafish larvae, either those subjected to TBI or uninjured controls, was recorded using Leica DM2500 LED optical microscope.

### Measuring the Seizure-like Phenotype in TBI Larvae

The seizure-like behavior and activity of zebrafish larvae post traumatic injury experiment was quantified via behavioral tracking software as described in our recent publications (Kanyo et al., 2020; Leighton et al., 2018). Briefly, control larvae or larvae subjected to traumatic brain injury using 65g weight, were placed individually in wells of 96-well plates. The locomotor and seizure activity were assessed 40 minutes after the traumatic brain injury through EthoVision® XT-11.5 software (Noldus, Wageningen, Netherlands). The hypermotility of larvae is a manifestation of Stage I and Stage II seizures (previously defined via application of epileptic drugs), whereas more intense Stage III seizures are arrhythmic convulsions that manifest as reduced macroscopic movement in this assay (Kanyo et al., 2020; Leighton et al., 2018; Liu and Baraban, 2019).

### Engineering CaMPARI transgenic zebrafish for integrative calcium imaging

The Tg[elavl3:CaMPARI (W391F+V398L)]^ua3144^ zebrafish line expressing the calcium sensor CaMPARI was generated using the Tol2 transgenesis system. We re-derived these previously established CaMPARI transgenic fish due to a federal moratorium on importing zebrafish into Canada (Hanwell et al., 2016). The Tol2 vector, pDestTol2-elavl3:CaMPARI (W391F+V398L), was a gift from Eric Schreiter’s lab and was published in (Fosque et al., 2015). The Tol2 vector was injected in embryos at the 1-2 cell stage as previously described (Fisher et al., 2006). Transient larvae were identified through green fluorescent cells in the central nervous system and were outcrossed to obtain a stable Tg[elavl3:CaMPARI (W391F+V398L)] ua3144 F1 line. The F1 line was crossed into the transparent Casper background.

### Measuring neuronal activity during TBI using CaMPARI

Bright green CaMPARI larvae were loaded into 20ml syringe containing 1ml E3 media (prepared as per Westerfield 2007, but without ethylene blue) and were exposed to a 405 nm LED array (Loctite), which illuminated the syringe entirely (Fig. 4A). Larvae were exposed for 10 sec, with the LED array at a distance of 7.5 cm from the syringe, while subjected to traumatic brain injury using the 300g weight as described above. Following this photoconversion of CaMPARI during TBI, larvae were anesthetized 0.24 in mg/mL tricaine (MS-222, Sigma Aldrich) and embedded in 2 % low-gelling agarose (A4018, Sigma Aldrich) for analysis under confocal microscopy.

**Fig 1.**
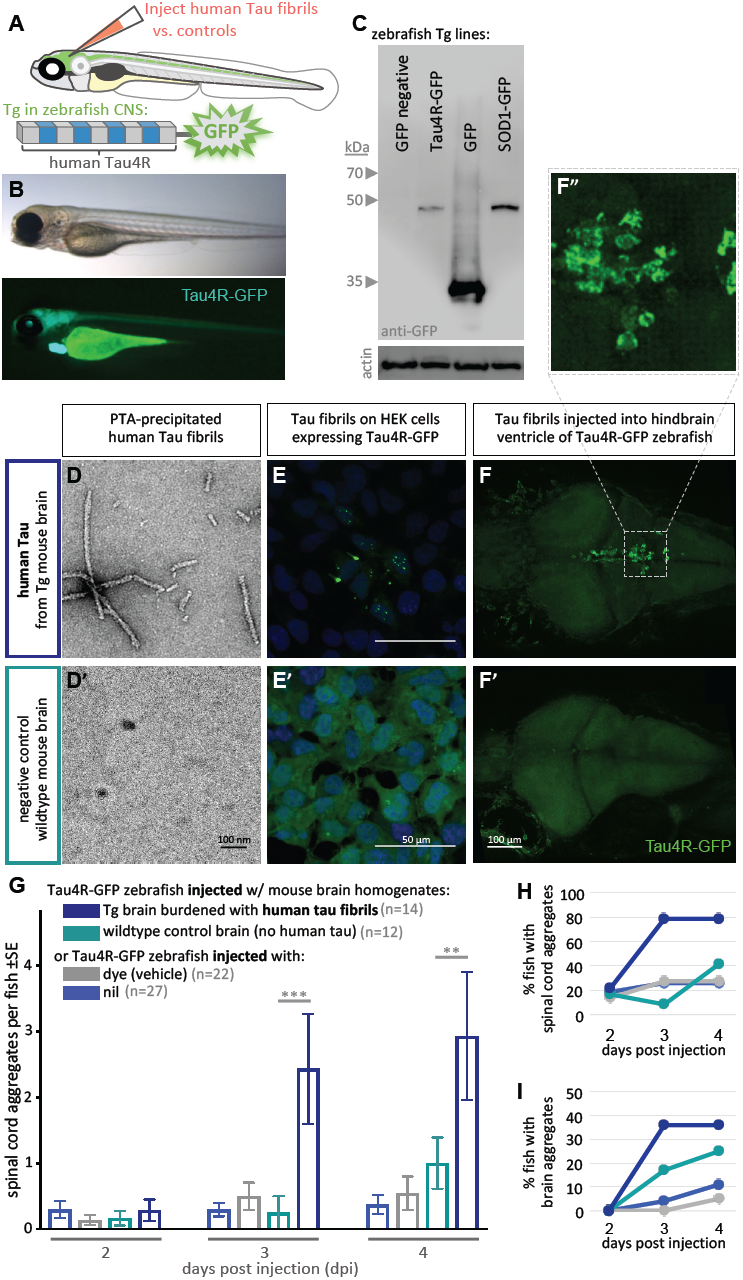
Validating tauopathy fluorescent biosensor *in vitro* and in zebrafish. The biosensor Tau4R-GFP was validated for its ability to detect tauopathy seeds *in vitro* and in zebrafish. **(A)** Schematic of Tau4R-GFP “Tau biosensor” that contains the four binding repeats (4R) region of wildtype human tau linked to green fluorescent protein (GFP; see also Fig S1A). **(B)** Transgenic zebrafish engineered to express Tau4 R -GFP biosensor throughout neurons of the CNS. Wildtype GFP is also abundant in the heart, which serves as a marker of the transgene being present but is otherwise irrelevant to our analyses. **(C)** Western Blot on zebrafish brain confirmed production of Tau4R-GFP at the expected size, similar to a SOD1-GFP biosensor and coordinately larger than GFP alone. **(D)** Human Tau fibril precipitated from transgenic (Tg TauP301L) mouse brain homogenates using PTA and assessed by EM. **(E)** Application of PTA-purified brain homogenate induced the formation of tau inclusions similar to clarified brain homogenate (scale bar 50μm; compare to Fig S1D), but application of equivalent preparations from non-Tg mice produced no GFP+ inclusions. **(F-I) Tau biosensor zebrafish detects diseaseassociated human tau fibrils following intraventricular injection of brain homogenate.** Crude brain homogenates were microinjected into the hindbrain ventricle of Tau4R-GFP zebrafish larvae at two days post-fertilization, and tau inclusions were analyzed at several time points. **(F)** Tau biosensor zebrafish larvae developed readily apparent GFP+ inclusions in the brain and spinal cord (Fig. S2) when injected with brain homogenate burdened with tau pathology (from Tg mice) but not from healthy brain homogenate (F’, from non-Tg mice). **F”** inset shows many adjacent cells exhibiting GFP+ Tau aggregates. **(G)** Tau biosensor zebrafish injected with human tau fibrils (within Tg mouse brain homogenate) developed significantly more aggregates on the spinal cord compared to uninjected control and other control groups, including compared to wildtype mouse brain homogenate (**p ≤ 0.01, ***p ≤ 0.001) **(H)** Same data as in G, expressed as the percentage of larval fish showing Tau aggregates in the spinal cord, and **(I)** those same fish also showed Tau aggregates in the brain, over time. n = number of individual larvae.

**Fig 2.**
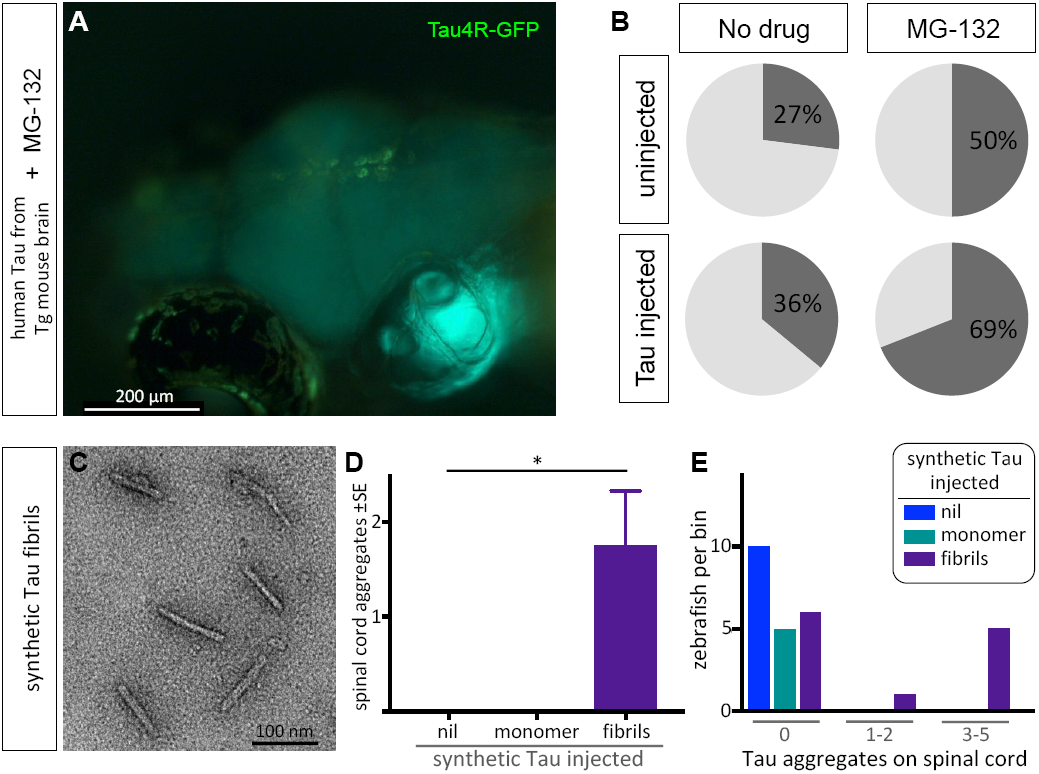
Protein-only induction of tau puncta *in vivo* detected in biosensor zebrafish. Injections of synthetic tau fibrils into Tau4R-GFP zebrafish induced GFP+ puncta in brains and spinal cord. **(A**,**B)** Inhibiting the proteosome with MG-132 enhanced the percentage of larvae bearing GFP+ inclusions in the brain following injection of tau-laden brain homogenate. **(C)** Synthetic human tau proteins were fibrillized as confirmed via EM analysis. Human Tau fibrils were microinjected into the larval hindbrain at two days post-fertilization, and tau inclusions were analyzed at three days post injections. **(D)** Tau aggregates were only observed after injection of tau fibrils, not monomers (*p<0.05). **(E)** Tau aggregates (same data as D, presented as distribution of larvae that displayed various amounts of GFP+ puncta) appear only after injection of tau fibrils, not monomers.

**Fig 3.**
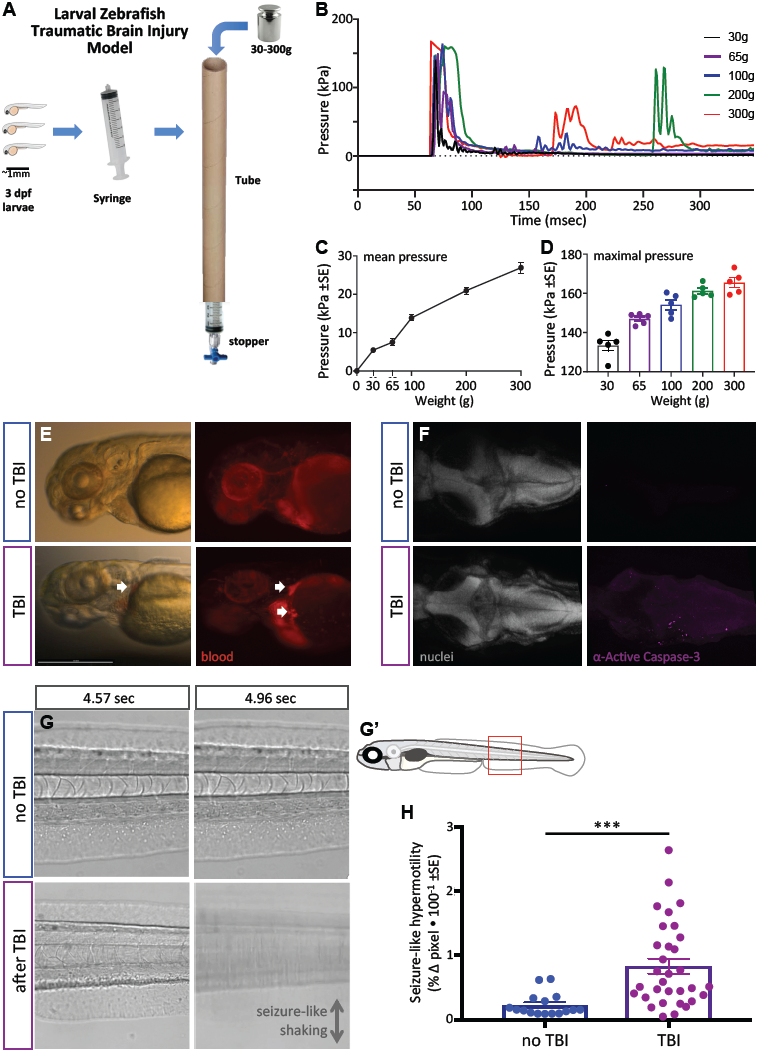
Zebrafish larvae subjected to traumatic brain injury (TBI) exhibited various biomarkers of TBI. **(A)** A novel TBI model for larval zebrafish: To induce blast injury, zebrafish larvae were loaded into a syringe with a stopper. A defined weight was dropped on the syringe plunger from a defined height, producing a pressure wave through the fish body akin to pressure waves experienced during human blast injury. **(B)** Dynamics of the pressure increase after dropping weights of varying masses in our TBI model. **(C**,**D)** The mean and maximum pressures generated, respectively, by various weights applied in the TBI model. Dots represent individual trials. **(E)** Hemorrhage after TBI was observed in some of the larvae fish using *Tg[gata1a:DsRed]* transgenic zebrafish that express DsRed in erythrocytes, as indicated by white arrows. Lateral view of larval heads with anterior at the left. **(F)** Increased cell death in the brain of 4 dpf larvae subjected to TBI as indicated by immunostaining of activated Caspase-3 (magenta). Positive and negative controls for immunostaining are in Figure S4. Nuclei were stained with DAPI in gray for reference. These are dorsal views of larval zebrafish brains with anterior at the left. **(G)** Seizurelike clonic shaking is observed in a subset of larvae after TBI. Movie frames are displayed from Supplemental Video S2. These frames (left and right panels) are separated by ∼400 msec in time, and are lateral views of the larval zebrafish trunk (akin to red box in G’). Control fish without TBI show little movement except obvious blood flow. Following TBI, larvae show bouts of calm (bottom left) interspersed (∼400 msec later) with bouts of intense seizure-like convulsions (Stage III seizures; bottom right). **(H)** Larvae subjected to TBI also displayed Stage II seizures, i.e. weaker seizures that manifest as hypermotility and are detected using a previously optimized behavioural tracking software system – seizures are significantly more intense following TBI compared to the control group (***p<0.001; dots are raw data for each larva, mean is plotted ±SE).

**Fig 4.**
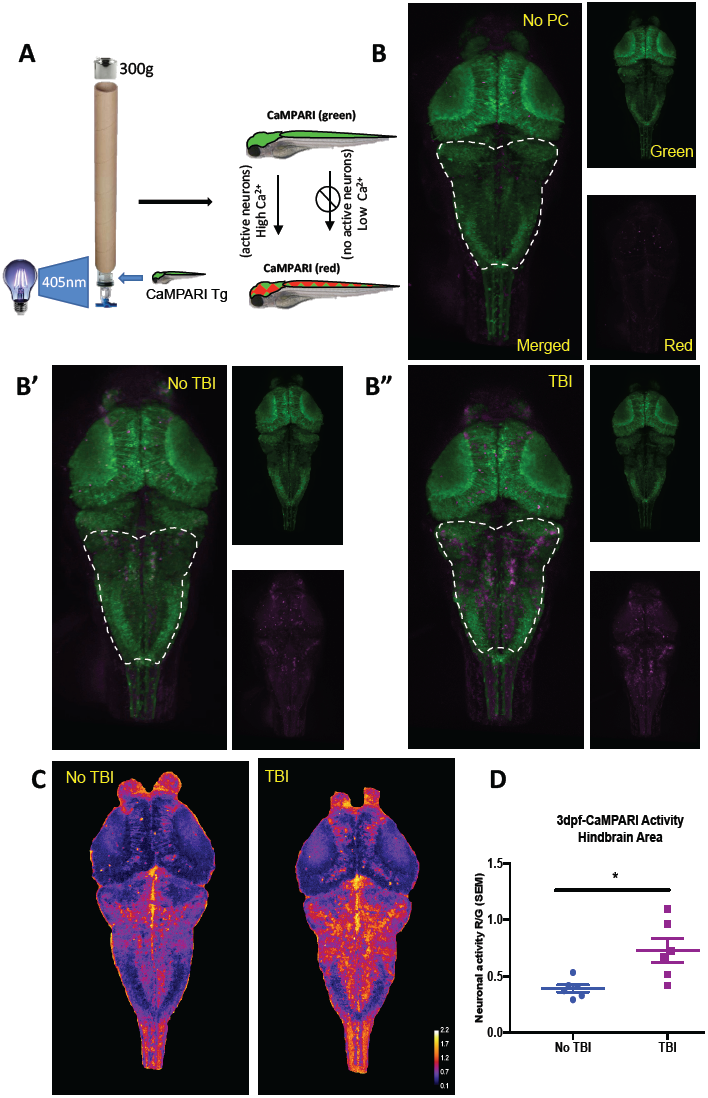
Neural activity increases during traumatic brain injury (TBI) as measured in CaMPARI zebrafish larva. **(A)** Schematic of TBI using CaMPARI (Calcium Modulated Photoactivatable Ratiometric Integrator) to optogenetically quantify neuronal excitability. 3dpf CaMPARI larvae were freely swimming while subjected to TBI, coincident with exposure to 405nm photoconversion light. CaMPARI fluorescence permanently photoconverts from green to red emission only if the photoconversion light is applied while neurons are active (high intracellular [Ca2+]). The ratio of red:green emission is stable such that it is quantifiable via subsequent microscopy. **(B)** Increased neural activity during TBI is represented by increased red:green emission (red pseudocoloured to magenta) in the hindbrain of larvae **(B”)**, compared to larvae not receiving TBI **(B’)** or fish not exposed to photoconverting light (“no PC” in panel B). These representative maximum intensity projection images show dorsal view of zebrafish brain (anterior at top), including merged, or red or green channels alone. **(C)** Heatmaps encode the CaMPARI signal (higher neural activity = higher red:green = hotter colours), highlighting location of increased neural activity during TBI relative to control larvae not receiving TBI. **(D)** Quantification of CaMPARI output in the hindbrain area reveals a significant increase in the neuronal excitability during TBI compared to control group not receiving TBI (*p<0.05. Each data point is an individual larva).

Maximum intensity projections were acquired from Z-stacks (8 µm steps) using a 20x/0.8 Objective and a laser-point scanning confocal microscope (Zeiss 700). The hindbrain area was analyzed, as it was the brain region most responsive to traumatic brain injury. To specifically isolate the brain regions and obtain data points, a 3D area was isolated by creating a surface with Imaris® 7.6 (Bitman, Zuerich) and the mean fluorescence intensities of the green and red channel intensities were calculated. Data points were presented as a red/green ratio for each individual larva and interpreted as relative neural activity, which is defined as red photoconverted CaMPARI in ratio to green CaMPARI (Fosque et al., 2015) (R Kanyo, IN REVISION Jan 16th, 2020).

### Bath Application of Drugs

Tau biosensor larvae were treated with 20 µM of the proteasome inhibitor MG-132 at 2dpf, following injections with brain homogenate from Tg human Tau mice. The treatment was left for 48 hours before changing the media and evaluating the percentage of larvae developing GFP+ puncta in the brain region.

For Kainic acid or kainate treatment (KA), the appropriate doses (5, 50, 100, 150, and 200μM) were added within 6 hours after TBI. For 4-aminopyridine (4-AP), one of two doses of 4-AP (200 or 800µM) were added either six or 24 hours after TBI, as indicated. For Retigabine (RTG) treatment, 10µM was used to treat TBI larvae beginning six hours after TBI. Unless otherwise stated, KA, 4-AP and/or RTG were applied to larvae for 38 hours, then a fresh drug-free E3 media was added. The formation of GFP-positive puncta was analyzed at four to five days post injury.

Pyrimidyn-7™(P7), the dynamin inhibitor, was purchased at a 50mM concentration supplied in DMSO (Abcam). Larvae that were subjected to TBI were treated within six hours following the injury with 3µM of P7. The dose was chosen based on the previous use of the P7 drug on zebrafish larvae (Verweij et al., 2019). The larvae were incubated with the drug for 20 hours, after which they were transferred to a fresh plate with drug-free media. Dyngo 4a, another dynamin inhibitor (McCluskey et al., 2013), was purchased from (Abcam) and 4 µM of Dyngo 4a was used to treat larvae as previously explained with P7. The formation and abundance of GFP-positive puncta was evaluated as previously described at four days post traumatic injury (dpti). For some experiments, the ‘Tau biosensor’ transgenic zebrafish were bred to a separate Tg line that express human four repeat TAU Tg(*eno2:hsa.MAPT-ires-egfp*)^Pt406^ throughout the zebrafish CNS (Bai et al., 2007).

### Statistics

All statistical analyses were performed using GraphPad Prism Software (Version 7, GraphPad, San Diego, CA). Sample sizes appropriate for our conclusions were estimated iteratively as the variance in each of our new methods became apparent; dose-response curves and significant differences amongst these dose were used to judge that any detected impacts of subsequent interventions would be valid. All experiments were independently replicated at least twice, individual larvae were the sampling unit (reported on Figures), and no outliers or other data were excluded. The experimenters were blinded to the treatments prior to quantifying outcomes. Unpaired t-tests were used to compare between two groups. For comparison between three or more groups at various time points or the same time point, 2-way and Ordinary One-Way ANOVA were used followed by post-hoc Mann-Whitney U tests and Kruskal-Wallace multiple comparison tests, respectively.

## Results

### Engineering and Validation of Tauopathy reporter lines

Previous reports describe the assessment and quantification of tau inclusions in living cells (typically Human Embryonic Kidney cells), via measuring aggregation of fluorescent proteins fused to tau protein, providing sensitive detection of pathological tau species and strain variants (Kaufman et al., 2016; Sanders et al., 2014; Woerman et al., 2016). Tau is predominantly expressed in the neurons of the CNS and we reasoned that fluorescent biosensor tools would have good potential to reveal additional phenotypes when expressed in these cells and moreover, that prion-like mechanisms of tauopathy spread are best modeled in an intact brain (e.g. vectored by blood and glymphatic circulation, ventricles, axonal projections and immune systems). Therefore, we engineered a tauopathy biosensor transgenic zebrafish that expresses a fluorescent tau reporter protein. Our genetically encoded fluorescent reporter protein was composed of the sequence of the human tau core-repeat domain fused to GFP with a linker sequence and is referred to here as Tau4R-GFP (Fig. 1A and S1A). Contrasting previous *in vitro* models, our biosensor did not feature any pro-aggregation mutations in the human tau repeats; this design was intended to minimize spontaneous aggregation events. The expression of the biosensor protein in zebrafish was under the control of the pan-neuronal promoter *neuronal enolase 2* (*eno2*, see Bai et al., 2007), which drives expression throughout the CNS (Fig. 1B and S1C). We deployed the transgene in a transparent zebrafish line (the ‘Casper’ background (White et al., 2008)) to facilitate analysis beyond the early larval development stages (when pigmentation would otherwise begin to obscure microscopy). We isolated a stable transgenic (Tg) line that expresses the Tau4R-GFP biosensor reporter robustly and clearly in the CNS (Fig. 1B), *Tg(eno2:Hsa.MAPT_Q244-E372-EGFP)*^*ua3171*^, and assigned it allele number ua3171.

Simultaneously, we expressed the same biosensor *in vitro* to validate the construct we deployed *in vivo* (Fig. S1A and B). Both in HEK293T cells and Tg zebrafish, immunoblotting using anti-GFP antibody detected our Tau-4R-GFP reporter protein at the expected size of ∼45 Kd, similar to a SOD1:GFP biosensor protein of similar predicted size, and an appropriately larger size relative to GFP protein alone (Fig. 1C, S1B’).

We assessed the capacity of our Tau4R-GFP biosensor to report the presence of tau pathology via transducing brain homogenates into cells. Brain homogenates burdened with tauopathy, from transgenic mice expressing mutant human tau (Tg Tau^P301L^), were compared to normal non-Tg mouse homogenates as a negative control. Congruent with findings obtained in past similar cell assays (Sanders et al., 2014), GFP-positive (GFP+) inclusions were detected only when cells were transduced with brain homogenate containing pathogenic human tau fibrils (from Tg Tau^P310L^ mice) (Fig. S1D). The *in vitro* assay detection rate was approximately 5% of cells having GFP+ inclusions in total, with 2% of cells forming multiple nuclear puncta and ∼3% forming one cytoplasmic inclusion, whereas various negative controls consistently displayed 0% of cells with inclusions (Fig. S1E). To verify that tau aggregates in the clarified brain homogenate caused the GFP+ puncta, we purified tau aggregates from the tissue samples using PTA precipitations (Woerman et al., 2016). Tau fibrils purified from these preparations were characterized via EM analysis (Fig. 1D). Transducing these preparations (in contrast to control preparations derived from non-Tg mice) produced fluorescent puncta in the Tau4R-GFP reporter cells (Fig. 1E), confirming the ability of our Tau4R-GFP chimeric protein to report tau aggregation.

### Validation of *in vivo* Tau biosensor via intra-ventricular brain injections of Tau fibrils

To test if the Tau4R-GFP biosensor can report the *in vivo* progression of tauopathy, we emulated intracerebral injection methods that induce (prion-like) tau pathology in mice (Clavaguera et al., 2013; Guo et al., 2016; Peeraer et al., 2015). We injected clarified brain homogenate laden with human tau fibrils, prepared as above from Tg mice, into the hindbrain ventricle of two days post-fertilization (dpf) tau biosensor zebrafish (Fig. S2A). The injected larvae and control groups were monitored daily for up to four days post-injection (dpi). Biosensor larvae injected with human Tau fibrils (from Tg mouse brain) developed GFP+ puncta, reflective of tau aggregation in the brain (Fig. 1F, F”). These tau inclusions were prominent near the ventricle wall as well as in sensory neurons along the spinal cord, when injected with brain homogenate from human-tau transgenic mouse (Fig. S2B). These puncta appeared to have either a lone dot-like shape or were similar to the multiple nuclear puncta detected *in vitro*, in which three to four small puncta are clustered together. Repeated assessment of the location of tau aggregates on the spinal cord of the same individuals over multiple days, using somite numbers as landmarks, suggested a movement of some of these puncta over time (Fig. S3A and B).

The abundance of GFP+ spinal cord inclusions was progressive and significantly higher in larvae injected with pathogenic TAU brain homogenate compared with various controls (p<0.0001 at 3dpi and 4dpi, Fig. 1G). Few larvae in the control groups developed spontaneous inclusions but the number of the larvae and the abundance of those inclusions were minimal (Fig. 1G). 80% and 35% of the larvae injected with human tau fibrils developed puncta in the brain and spinal cord, respectively (Fig. 2H and I). On the other hand, a lower proportion of tau biosensor larvae developed ‘spontaneous’ inclusions post-injection with the control brain homogenate from wild-type mice (Fig. 2H and I). Tau aggregates were detected on the spinal cord region as early as 2 dpi. Intriguingly, a small percentage of larvae developed sporadic GFP+ tau aggregates regardless of treatment. Visualizing the data as distributions of larvae with particular abundances of GFP+ inclusions (Fig. S2C) highlights a trend where most larvae did not develop aggregates unless they were injected with brain homogenate containing fibrillar, pathogenic human tau species. In those cases, the biosensor larvae developed an abundant number of aggregates. Overall, these data confirm the ability of our biosensor model to detect pathogenic tau species *in vivo*.

Like other protein misfolding diseases, tauopathies reflect a proteostatic imbalance wherein the clearance of pathological tau species is insufficient relative to accumulation (Chiti and Dobson, 2006; Lim and Yue, 2015). We reasoned that if the tau biosensor larvae are faithfully reflecting tau proteostasis concepts *in vivo*, then this could be revealed via inhibition of the proteasome. We treated larvae with the proteasome inhibitor MG-132 (Fig. 2A) and observed an approximate doubling of spontaneous GFP+ inclusions (Fig. 2B). Following injection of mouse brain homogenate containing human tau fibrils, applying the proteasome inhibitor MG-132 substantially enhanced the percentage of larvae bearing Tau4R-GFP+ inclusions in the brain (to ∼70%, Fig. 2B), relative to equivalent larvae without MG-132 (∼36%, Fig. 2B).

It was striking that the zebrafish tau biosensor was robustly able to discriminate brain homogenates with human tau aggregates versus those that were not. However, we considered an alternative explanation for the data: the difference may not depend directly on human tau in the brain homogenate but could instead reflect other bioactive components of the degenerating Tg mouse brain. To verify that the formation of GFP+ puncta in zebrafish can be seeded by a protein-only injection, we delivered synthetic human tau protein (2N4R). After confirming the recombinant tau proteins were appropriately fibrillized via EM (Fig. 2C), we delivered them by intraventricular injections as described above. Similar to previous data with brain homogenate, the larvae that were injected with synthetic tau fibrils developed inclusions proximal to the brain ventricles as well as along the spinal cord at 3-6 dpi. The abundance of tau aggregates along the spinal cord was significantly higher in larvae injected with the synthetic tau fibrils compared to larvae injected with tau monomers or to the non-injected group (p<0.05) (Fig. 2D). The distribution of larvae based on the number of tau aggregates they accumulated also supported these findings (Fig. 2E). In sum, the Tau4R biosensor deployed in the CNS of larval zebrafish was able to report tau species, and further revealed the prion-like induction of tauopathy via protein-only seeding *in vivo*.

### Introduction of the first traumatic brain injury model for larval zebrafish

We next sought to deploy our tau biosensor in a tauopathy model that enables higher throughput than can be achieved with intraventricular injection methods. We considered traumatic brain injury (TBI) as an inducer of the tauopathy in Chronic Traumatic Encephalopathy (CTE); further, we were encouraged that innovations in this realm could fill an unmet need for a high-throughput, genetically tractable *in vivo* model of these devastating concussive injuries. Although a few methods have been reported to induce traumatic brain injury in <u>adult</u> zebrafish that are comparable to mammalian TBI methods (Maheras et al., 2018; McCutcheon et al., 2017), no such methods were available for zebrafish larvae. Here we introduce and validate a simple and inexpensive method to induce traumatic brain injury in zebrafish larvae. Investigating traumatic brain injury in larvae offers substantial benefits regarding experimental throughput, economy, accessibility of drug and genetic interventions, and bioethics.

We devised a traumatic injury paradigm by loading zebrafish larvae (∼12 individuals in their typical E3 liquid growth media) into a syringe with a closed valve stopper, and applying a hit on the plunger to produce a pressure wave through the fish body akin to pressure or shock waves experienced during human blast injury (Nakagawa et al., 2011) (Fig. 3A). To challenge the method’s reproducibility, and to permit manipulation of injury intensity, a series of defined masses were dropped on the syringe plunger. Technical variability, anticipated from larvae being in different orientations and positions within the syringe, was reduced by applying the injury three times to each group of larvae (except where noted otherwise) while repositioning the syringe between each injury. To assess if our method faithfully induced traumatic brain injury similar to injury from pressure waves, we examined multiple markers known to be associated with blast-induced traumatic brain injury, including cell death, hemorrhage, blood flow abnormalities, and tauopathy (Bir et al., 2012; Kovacs et al., 2014; Nakagawa et al., 2011). Additionally, we evaluated the occurrence of post-traumatic seizure activity and increases in neuronal activity acutely associated with the trauma.

We established the TBI method via empirical testing of various parameters, restricting ourselves to materials and methods that can be adopted inexpensively, with a goal of consistently inducing a robust injury (see phenotypes below) vs. a tradeoff with maximizing survival of the larvae. Subsequent to this optimization, we were able to characterize the pressure induced within the syringe during each injury (Fig. 3B-D). The maximum pressure induced was near 170 kPa (Fig. 3B). The dynamics of the pressure change events during TBI (Fig. 3B) imply that dropping the heavier weights led to the weight bouncing and producing a secondary increase in pressure (e.g. at ∼175 or ∼275 msec in Fig. 3B). The maximal pressure induced varied from ∼130 to ∼175 kPa in an approximately linear fashion depending on the mass of the weight dropped (Fig. 3D). The mean pressure change over the first 300 msec of the TBI also increased in a nearly linear fashion, and increased by nearly an order of magnitude when dropping weights of 30g compared to 300g (Fig. 3C).

We evaluated TBI-induced hemorrhage via the use of *Tg[gata1a:DsRed]* larvae that have red fluorescence in their blood cells (Traver et al., 2003). Hemorrhage was observed variably in larvae when a heavy weight (300g) was used to induce the traumatic injury (Fig. 3E). Further, approximately half of the TBI larvae showed abnormalities in blood flow including a temporary reduction or complete absence of blood circulation (video 1), consistent with abnormalities detected in rodent TBI models (Bir et al., 2012). Subsequently, we assessed apoptosis in the TBI larvae, observing that our TBI method induced cell death in larvae as detected by staining for active Caspase-3 (Fig. 3F and S4). The number of active-Caspase-3-positive cells was negligible in the control groups compared to a mean of 62 apoptotic cells in TBI larvae (SEM ± 9.17, n=3) and 75 (SEM ± 4, n=2) in positive-control-larvae (cell death induced with camptothecin, CPT; Fig. S4). These data all align well with existing animal models of TBI with respect to mimicking characteristic features of human TBI, and support the effectiveness of our method in inducing traumatic brain injury in larval zebrafish.

### TBI treated larvae exhibited post-traumatic seizure-like behavior and increased neuronal activity during Trauma

Post-traumatic seizures are one of the most frequent conditions associated with traumatic brain injuries and, despite being prevalent, remain poorly understood in TBI patients (Kovacs et al., 2014). Post-traumatic seizures were overtly apparent in a subset (approximately 40%) of zebrafish larvae after they were subjected to traumatic brain injury. In some instances the activity was highly reminiscent of Stage III seizures (defined previously in larval zebrafish as the most intense seizures; Liu and Baraban, 2019) with bouts of intense clonic convulsions and arrhythmic shaking (Supplemental Video S2; exemplar frames from the movie are in Figure 3G). Other individuals exhibited hypermotility that is exactly consistent with past definitions of less intense Stage I or Stage II seizures. We quantified the latter seizure activity via behavioral tracking software (which we had previously optimized and validated for quantifying seizures in larval zebrafish (Kanyo et al., 2020; Leighton et al., 2018)) and determined that larvae subjected to TBI exhibited seizure-like activity that was significantly higher than the control group (p<0.0007) (Fig. 3H).

Seizures are caused by abnormal and excessive neuronal excitability (Stafstrom and Carmant, 2015). To document bursts of neuronal activity *during* the brain trauma, if any, we utilized a genetically encoded calcium imaging CaMPARI reporter (calcium modulated photoactivatable ratiometric integrator) expressed throughout the CNS. CaMPARI fluoresces green in baseline conditions, and permanently converts to red fluorescent emission if high intracellular calcium levels (i.e. neural activity) occur coincident with application of ‘photoconverting’ intense 405nm blue light. We subjected our allele of *Tg[elavl3:CaMPARI]*^*ua3144*^ larvae (R Kanyo, IN REVISION Jan 16th, 2020) to TBI, coincident with brief application of photoconverting light (405nm light provided by an LED array directed at the syringe, as described in Fig. 4A). A sharp increase in neuronal activity during TBI was evident, especially in the hindbrain region as indicated by enhanced red emission (Fig. 4B). CaMPARI allows robust quantification of neural activity expressed as a ratio of red:green fluorescent emission, which confirmed that neuronal excitability increases significantly in response to brain trauma (Fig. 4C and D). Notably, this combination of newly introduced methods of traumatic brain injury being integrated with CaMPARI optogenetic methods (where the stable/irreversible changes from green to red fluorescent reportage allows a ratiometric quantification in a subsequent microscopy session) offers the rare ability to assess neural activity on unrestrained (free-swimming) subjects during TBI. In sum, our data reveal a substantial burst of neural activity occurs *during* TBI, and that zebrafish larvae exposed to TBI subsequently exhibit a significantly higher propensity for spontaneous seizures.

### Traumatic brain injury on Tau biosensor zebrafish larvae induced GFP+ puncta

After validating that our method was able to induce traumatic brain injury upon zebrafish larvae, we next asked whether TBI induces tau aggregates in our tau biosensor model. Initially, we evaluated if our TBI method would induce aggregation of fluorescent proteins in models expressing GFP alone or other biosensor proteins such as SOD1-GFP. Following TBI, and regardless of injury intensity, no GFP+ aggregates were detected in these controls (Fig. S5 A-C). Similar results were obtained with other transgenic zebrafish that express GFP in motor neurons (data not shown). Further, our Tau4R-GFP fish additionally express an unmodified GFP variant in the active heart muscle, and this robust GFP showed no sign of aggregation following TBI. Remarkably, in these same individual Tau4R-GFP larvae we detected Tau4R-GFP biosensor GFP+ puncta in both brains and spinal cords following TBI (Fig. 5A-B). The abundance of GFP+ puncta increased with time following the injury (Fig 5C-D). To determine if the severity of tauopathy varies coordinately with severity of the traumatic injury, we assessed the impact of different masses. Although some variability is evident, a dose-response relationship is apparent such that the 65g, 100g and 300g weights induced more tau aggregates compared to the control and 30g weight (Fig. S8E). The heaviest weight (300g) induced significantly more tau aggregates versus the control group or the group with the 30g weight (p<0.01 and p<0.001, respectively). Therefore, we decided to use both the 300 and 65 g weights for subsequent experiments. We evaluated whether dropping the light weight once or multiple times would affect the number of tau aggregates on the spinal cord as well as dropping the weight once on three consecutive days, perhaps reminiscent of repetitive sports injury. We observed an increase in the abundance of tau aggregates when the weight was dropped multiple times during one day, or over three consecutive days, but this increase was not statistically significant (Fig. S8 B-D).

**Fig 5.**
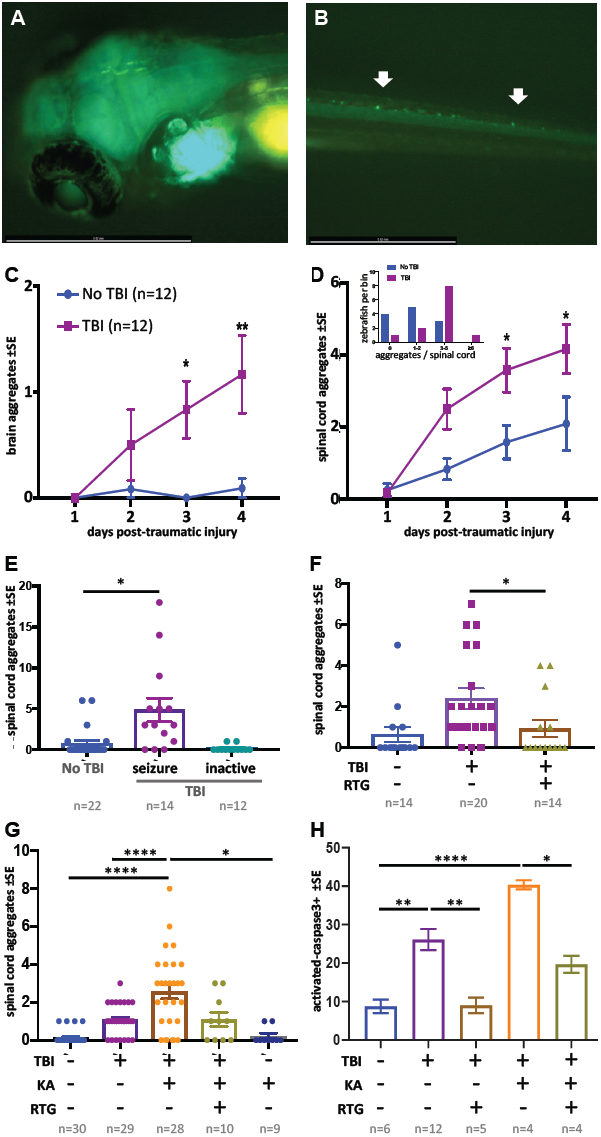
Traumatic brain injury (TBI) induces tauopathy in larval zebrafish. **(A)** GFP+ Tau puncta are detected in the brain of Tau4R-GFP biosensor zebrafish at 5 days post traumatic brain injury (dpti). **(B)** Tau aggregates formed on the spinal cord as a result of the traumatic brain injury as shown by arrows. **(C)** Tauopathy significantly increases over time following TBI compared to control group (No TBI) (*p<0.05 **8** at 3dpti and **p<0.01 at 4dpti). **(D)** The number of tau aggregates in spinal cord significantly increases over time following TBI compared to control group (*p<0.05 at 3dpti and 4dpti). Inset: Tau4R-GFP zebrafish larvae subjected to TBI develop more GFP+ puncta compared to the control group by 3dpti (inset plot is similar to Fig. 2E). **(E-H) Post-traumatic seizures link TBI to tauopathy. (E)** Following TBI, larvae displaying post-traumatic seizures developed many more tau aggregates relative to those not displaying post-traumatic seizures (***p<0.001). **(F)** Inhibiting post-traumatic seizures with the anti-convulsant retigabine (RTG, 10μM) significantly decreased the abundance of GFP+ puncta in the spinal cord (p<0.05). **(G)** Increasing post-traumatic seizure using the convulsant kainate (KA, 100μM) significantly increased the formation of tau aggregation following TBI; this effect was prevented by co-treatment with anti-convulsant RTG. **(H)** Blunting post-traumatic seizures with RTG reduced TBI-related cell death. The main impact of RTG was specific to its anticonvulsant modulation of seizures because its effects were reversed by convulsant KA. Colour scheme in panel C applies to other panels. n= number of zebrafish larvae.

The GFP+ tau aggregates formed in the brain region following TBI tend to form fused shapes (Fig. S8A) reminiscent of the spontaneous aggregates described above. The aggregates on the spinal cord, however, had similar shapes to aggregates detected post brain-injections, but with qualitatively less brightness in some instances. Multi-day monitoring of individual larvae (Fig. S6 & S7) revealed variation in formation of tau aggregates amongst individual TBI larvae. We monitored the abundance of tau aggregates within individual fish over time following TBI and found that the average tauopathy significantly increased compared to the control group (p<0.05 at 3 dpti and p<0.01 at 4 dpti (days post traumatic injury) (Fig. 5C,D). Analysis of distribution of larvae binned into the number of Tau4R-GFP+ puncta at 3 dpti showed that more larvae developed Tau4R-GFP+ puncta compared to the control group (inset in Fig 5D). Considering that many of the larvae subjected to TBI formed Tau4R-GFP+ puncta in the brain that had a fused pattern (Fig. S8A), we focused on tau aggregates that formed on the spinal cord as their abundance could be most efficiently quantified compared to aggregates that formed in the brain.

### Post-traumatic seizure intensity influences tauopathy progression

Considering the clinical prominence of post-traumatic seizures following TBI, and the suggested role of cell stress and increased neural activity in promoting protein misfolding diseases (Kovacs et al., 2014; Sanchez et al., 2018), we speculated that post-traumatic seizures might form a causal link between TBI and subsequent tauopathy. We first asked if a correlation exists between seizure intensity and extent of tauopathy. Following TBI, some larvae exhibited seizure-like movements, while some did not seem to move abnormally relative to untreated fish (Fig. 3H). We sorted the larvae subjected to TBI into groups exhibiting the seizure-like behavior and those that displayed no overtly abnormal movement. Larvae exhibiting seizure-like behavior after TBI went on to develop abundant spinal cord aggregates (5-fold increase, p<0.001) in comparison to larvae that showed no seizure-like response to TBI (Fig. 5E).

To assess the hypothesis that seizure activity has a causal role in increasing the abundance of tau aggregates in our TBI model, we employed convulsant and anti-convulsant drugs to modulate the seizure intensity and *in vivo* neural activity. We selected drugs that are well-established to behave similarly in zebrafish as in mammals, though it is perhaps notable that the multi-day drug application used here is longer than the acute applications typically considered in zebrafish (Ellis et al., 2012). Our hypothesis predicted that decreasing seizure-like activity following TBI would reduce tauopathy. Indeed, applying the anti-convulsant drug Retigabine (RTG), that opens voltage-gated potassium channels (KCNQ, Kv7), resulted in a significant decrease in the abundance of GFP+ puncta (p<0.05) with many TBI larvae not developing any Tau4R-GFP aggregates (Fig 5F).

Similarly, intensifying post-traumatic seizures via application of the convulsant kainate increased the abundance on tauopathy 4-fold (p<0.001. Fig 5G) in a dose-dependent manner (Fig S9). Kainate did not increase Tau4R-GFP+ puncta in the absence of TBI. Surprisingly, the convulsant 4-aminopyridine did not increase tauopathy (explored below).

To assess if the impacts of kainate and retigabine on tauopathy were directly due to their modulation of post-traumatic seizures, we applied effective doses of each in concert. Co-application of kainate and retigabine following TBI produced an abundance of Tau4R-GFP+ puncta that was indistinguishable from larvae receiving TBI without pharmacology (Fig 5G).

TBI-induced cell death was likewise correlated with the intensity of post-traumatic seizures. Co-application of kainate and retigabine following TBI increased or decreased, respectively, the abundance of cell death in a manner coordinate with the tauopathy (Fig 5H).

Overall, convulsant and anti-convulsant drugs acted to increase and decrease TBI-induced tauopathy, respectively. The drugs appear to be specific – their individual impacts on tauopathy and cell death are largely attributable to their epileptic and anti-epileptic modulation of post-traumatic seizures, because when kainate and retigabine were applied concurrently they negated each others’ effects.

### Appearance of tauopathy following TBI requires endocytosis

To further examine increased seizure activity after TBI, we applied 4-aminopyridine (4-AP), a K_v_ channel blocker and convulsant drug. We predicted that raising the level of seizure activity would elevate tauopathy abundance in our TBI model, aligning with our observations following application of kainate (above). Surprisingly, higher doses of 4-AP consistently abrogated the appearance of tau aggregates. Treating TBI larvae with 200 or 800μM of 4-AP for a prolonged period (38 hours, beginning 24 hours post traumatic injury) significantly inhibited the abundance of Tau4R-GFP+ puncta in the TBI group (Fig. 6A-B, Fig. S10A-B). Analysis of the distribution of larvae linked to the number of tau aggregates supported this finding with no zebrafish larvae developing aggregates in groups treated with 4-AP (Fig. S10C). It is worth noting that 4-AP is commonly used in zebrafish models of epilepsy, but rarely used for prolonged treatment. To evaluate if the time at which treatments are administered plays a role in this unexpected result, we treated larvae with 200 μM 4-AP at earlier time points, specifically during traumatic brain injury and 1.5 hours later. We kept the duration of 4-AP treatment the same as previous experiments (38 hours). We found that administering 4-AP during different time windows relative to the traumatic brain injury did not measurably alter the inhibitory action of 4-AP on the abundance of tau aggregates (Fig. S10D). A similar observation was made when the duration of the 4-AP treatment was reduced to 24 hours (Fig. S10E).

**Fig 6.**
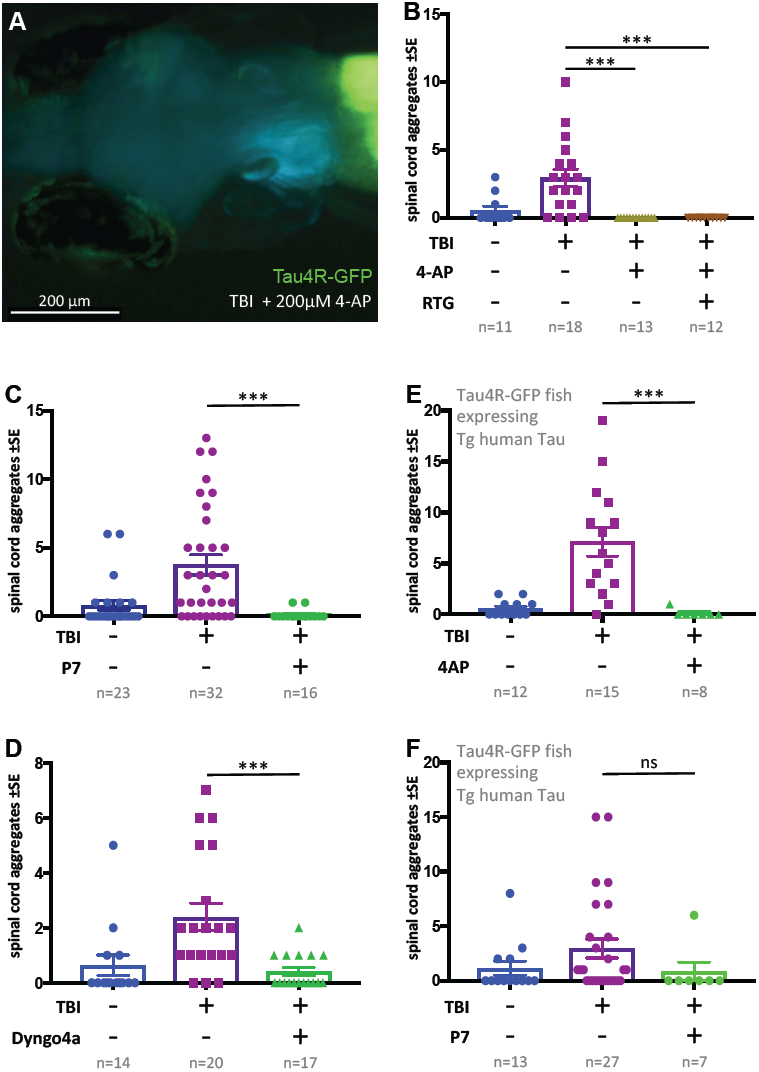
Tauopathy induced by traumatic brain injury (TBI) was attenuated by 4-aminopyridine (4-AP) via mechanisms independent of seizures. **(A)** Tau4R-GFP biosensor zebrafish larvae subjected to TBI and treated with the convulsant 4-AP show no brain puncta. **(B)** 4-AP significantly reduced (apparently eliminated) the abundance of GFP+ puncta in the brain and spinal cord compared to untreated TBI control. The impact of 4-AP on tauopathy appears to be independent of its actions on post-traumatic seizures because reducing the latter with anti-convulsant retigabine (RTG) had no measurable effect. See also Figure S10. **C-F. Pharmacological inhibition of endocytosis reduced tauopathy following TBI. (C)** Blocking endocytosis with Pyrimidyn-7 (P7) treatments significantly inhibited the formation of Tau4R-GFP+ puncta following TBI in zebrafish larvae (***p<0.001). **(D)** Dyngo 4a treatment significantly reduced Tau aggregates in the spinal cord (***p<0.001) in a manner similar to P7. **(E)** 4-AP treatment significantly inhibited the formation of Tau4R-GFP+ puncta in the spinal cord (***p<0.001) of Tau biosensor line that also express human Tau (0N4R) after traumatic brain injury compared to untreated TBI control group. **(F)** A notable reduction in Tau aggregates was observed in the same line after treatment with P7 drug. Statistical analysis shows no significance difference between groups. n= number of larvae.

Next, we considered if this unexpected inhibition of tauopathy by high-dose 4-AP convulsant is a direct consequence of increased neural activity (e.g. perhaps via neural exhaustion). We found that larvae receiving TBI and 4-AP continued to exhibit a lack of tau aggregates when co-treated with anti-convulsant retigabine (p<0.0001) (Fig. 6B). This suggested that high doses of 4-AP block the formation of tau aggregates via a mechanism independent of its convulsant activity.

To resolve a mechanism whereby high doses of 4-AP reduced tau pathology, contrary to our predictions above regarding neural hyperactivity, we considered previous *in vitro* work that demonstrated high concentrations of 4-AP cause reduced endocytosis of synaptic vesicles (Cousin and Robinson, 2000). To examine if the inhibitory actions of 4-AP on the abundance of tau aggregates in our TBI model is consistent with an endocytosis inhibition mechanism, we treated our tau biosensor larvae post traumatic injury with Pyrimidyn-7 (P7), a potent dynamin inhibitor that is known to block endocytosis (McGeachie et al., 2013), and analyzed the propagation of tau pathology by quantifying the number of tau inclusions. Owing to the potency of P7 and its impact on the survival of larvae, we treated the larvae with it for 24 hours at 3 μM. Similar to the findings with 4-AP, P7 treatments significantly inhibited the formation of Tau4R-GFP+ puncta in TBI larvae (p<0.001) (Fig. 6C). We assessed further the role of endocytosis by employing another dynamin-inhibitor drug, Dyngo 4a, that is less potent than P7 (McCluskey et al., 2013). We obtained similar results in which Dyngo 4a treatments significantly reduced tauopathy in our TBI model (Fig. 6D). To determine if these results are applicable to human tau, we induced traumatic brain injury on double-transgenic larvae expressing both human tau (the 0N4R human Tau isoform, see Bai et al., 2007) and our tau biosensor reporter, followed by treatment with either 4-AP or P7. Apart from the untreated control, both groups treated with 4-AP or P7 exhibited a noticeable reduction in abundance of tau aggregates. While the decrease in the case of P7 was not statistically significant, statistical analysis showed significance after 4-AP treatments (p<0.001) (Fig 6E and F). These findings confirmed the ability of 4-AP and dynamin inhibitors of reducing human tau aggregates in our TBI larvae.

## Discussion

The consequences of concussive blasts and TBI extend beyond the proximate injury – they are prominent risk factors for devastating dementias including AD, CTE and other tauopathies. Identifying the causal links that entwine TBI with subsequent tauopathies would inspire improved diagnostics and therapeutics. Investigating these mechanisms has been hampered by lack of tractable models, since cell culture platforms cannot faithfully represent the injury or the response-to-injury or the treatment thereof. Indeed, TBI and tauopathies are complex tissue and systems-level events with pathobiology progressing on a backdrop of dynamic vigorous neural function, prion-like vectoring of misfolded proteins via glymphatic and blood vasculature, immune and support cells, sleep physiology, homeostatic regulation and complex drug metabolism. Rats have been the favoured animal model for TBI, and mice can complement this as insightful models of tauopathy, yet both are challenged by expense, ethical considerations, and CNS tissues that are relatively inaccessible to (longitudinal) visualization of cellular events in living individuals (Bodnar et al., 2019; Marklund, 2016; Meconi et al., 2018; Pham et al., 2019). Here, our zebrafish models imperfectly replicate human TBI and tauopathy, but by addressing many of these challenges in an accessible and vibrantly active vertebrate brain, we offer an innovative approach for the study of prion-like events, tauopathy and/or TBI.

Here, we introduce a simple method for delivering TBI to larval zebrafish that can be scaled to high-throughput and adopted at low expense. The tractability and transparency of zebrafish larvae allowed us to deploy genetically encoded fluorescent reporters that were validated to (i) uniquely quantify neural activity on freely behaving animals *during* TBI, and (ii) effectively document prion-like tauopathy in individual subjects over multiple days. The accessibility of this platform to pharmacology allowed us to query cell biology events *in vivo* and support a role for endocytosis in prion-like progression and TBI-induced tauopathy. Further, anti-convulsant drugs were potent mitigators of the tauopathy and cell death that emerged subsequent to TBI; these effects were attributable to suppression of post-traumatic seizures (as proven by the anticonvulsant’s therapeutic effects being reversed by co-application of convulsant drugs). It remains to be seen if this data has any bearing on the long-term clinical management of TBI patients, and we newly speculate that prophylactic application of anti-convulsants (already common for blunting of patient’s post-traumatic seizures) might hinder progression of tauopathies including CTE and AD. If true, then debates regarding the optimal regimen of anti-epileptics for TBI patients should consider their potential for providing long-term benefits on dementias.

### Prion-like tauopathy induced by TBI

It is well established that TBI induces tauopathy, and over the past decade, numerous *in vitro* and *in vivo* studies have supported that tau proteins possess prion-like properties. Indeed seeding, templated misfolding (conversion) and spread to synaptically connected regions has been documented in tauopathies such as AD and FTD (Ayers et al., 2018; de Calignon et al., 2012; Goedert et al., 2017a; Goedert et al., 2017b; Iba et al., 2015; Woerman et al., 2016). Various mechanisms have been proposed for the transcellular transfer of tau seeds including release mechanisms via exosomes, or cellular uptake mechanisms via endocytosis (Demaegd et al., 2018; Evans et al., 2018; Wang et al., 2017; Wu et al., 2013). Nonetheless, these suggested mechanisms were postulated based on *in vitro* evidence as there is a lack of appropriate models that can visualize and manipulate the prion-like spread of tau pathology between tissues in a vibrant brain. It is unclear if these mechanisms are universal in progression of all tauopathies or if there are factors and mechanisms that are unique to each disease. Our zebrafish models allow us to study the progression and spread of TBI-induced tauopathy longitudinally in living animals, an experimental advantage that is unmatched among TBI animal models.

Regarding TBI, a large knowledge gap exists regarding how tau seeds are released and/or internalized by adjacent (or far-flung) cells - indeed the prion-like properties of tau species following TBI had not been assessed until very recently (Woerman et al., 2016; Zanier et al., 2018). Moreover, the focus in the literature has mostly been directed towards repetitive mild trauma as it is most associated with CTE, yet the various forms of TBI all are considered risk factors for neurodegeneration. The recent revelation that most TBI patients, whether they suffered from single or repetitive brain trauma, all exhibited tau pathology similar to CTE (Washington et al., 2016; Zanier et al., 2018) suggests all forms of TBI might incorporate tauopathies. Our mode of TBI on larval zebrafish entails a pressure wave that most closely mimics a blast injury (e.g. as experienced by military personnel or civilians near an explosion), but the etiology leading to tauopathy probably has many similarities regardless of the mode of the initiating TBI.

### Post-traumatic seizures accelerate tauopathy following TBI

We inspected the role of seizure activity and/or neuronal excitability, as well as the role of dynamin-dependent endocytosis, during the progression of tauopathy after TBI. We focussed our attention on seizure activity in part because seizures frequently occur in TBI patients following blast traumatic injury (Englander et al., 2014; Kovacs et al., 2014). We hypothesized that neuronal excitability and seizure activity after TBI can play a role in accelerating the wide dissemination of tau pathology. As such, we introduced two new approaches to test this hypothesis. The first approach was to engineer a novel *in vivo* tau biosensor model in zebrafish that can visualize pathological tau spreading and accumulation within the intact and vibrant CNS. The tau biosensor zebrafish express human tau4R-GFP reporter protein, and we confirmed its ability to detect tau seeds from various sources both *in vivo* and *in vitro*, similar to previously engineered *in vitro* models (Kaufman et al., 2016; Sanders et al., 2014). Our second approach was to introduce and optimize an elegantly simple technique to cause pressure-wave induced TBI, similar to human blast TBI, in zebrafish larvae. We endeavoured to inflict injury on larval zebrafish rather than adults because of the synergistic advantages that larval zebrafish provide: these include economical access to large numbers of individuals and associated statistical power, and the tractability of larvae for high-throughput *in vivo* screening of therapeutic agents (Saleem and Kannan, 2018). Larval zebrafish provide a large economic advantage compared to adults, with respect to time, cost per individual and space consumed in animal housing. Moreover, injuring animals in larval stages is viewed as an ethically favourable Replacement [*sensu* “the three Rs” of (Russell and Burch, 1959)] compared to injuring adult subjects. Thus, regardless of any bioethical considerations based on taxonomy, larval fish (that are accessible early in their development via external fertilization of eggs) are ethically advantageous to rodents (that are accessible for TBI only at postnatal stages) when considering highly invasive procedures like TBI. The latter conclusion relies on the assumption that the knowledge gained is of value, i.e. relevant to appreciating disease etiology.

Our data argue that our TBI methods are germane to clinical aetiology, because (akin to existing animal models of TBI) we were able to confirm the presence of various markers associated with brain injury, such as cell death, abnormalities in blood flow, hemorrhage and the occurrence of post-traumatic seizures.

The post-traumatic seizures apparent in our TBI model led us to consider the neural events occurring during the TBI, and their potential bearing on the correlation between neural activity and tauopathy. Few studies examine how TBI impacts neuronal circuits, especially *in vivo*, and these typically consider events several hours or days after brain trauma (Bugay et al., 2019). This may be of importance when considering evaluating the reasons behind the developments of post-traumatic seizures and epilepsy. In a controlled cortical impact model of TBI, an initial decrease or loss in neuronal activity is recorded after injury before a rise in neuronal activity is noted (Ping and Jin, 2016). Whether this occurs in different types of TBI, like blast TBI, was unexamined. To address this, we performed TBI on larval zebrafish expressing CaMPARI, a genetically encoded optogenetic reporter of neural activity. CaMPARI is particularly ideal for this question, as its reportage of neural activity (a stable and quantifiable shift from green to red fluorescence) occurs only during user-defined times and that reportage is relatively permanent. This allowed us to quantify the CNS activity that had occurred during TBI, by characterizing the ratio of red:green fluorescent emission using confocal microscopy after the TBI injury was completed. This approach therefor allows relatively easy access to quantifying neural activity *during* injury in an unencumbered freely-swimming animal. Here, we revealed for the first time a snapshot of neurons becoming active at the moment of TBI. Our results demonstrated an increase in neuronal excitability upon TBI, which may contribute to the frequency of post-traumatic seizures observed in our model, other blast TBI models and TBI patients (Bugay et al., 2019). Regarding the etiology of tauopathy subsequent to TBI, the CaMPARI quantification provided us important validation that neural activity was substantively impacted by TBI, complementing the evidence of increased seizure-like movements. This supported our rationale that convulsant and anti-convulsant drug treatments might modulate neural activity and thereby accelerate or decelerate tauopathy accumulation.

Indeed we had noted the occurrence of post-traumatic seizures in most of our TBI samples, which is in agreement with the prevalence of seizures in blast TBI patients and TBI rodent models (Bugay et al., 2019; Kovacs et al., 2014). However, whether post-traumatic seizures contribute to prion-like spreading of tau pathology (observed after TBI or not) was unknown. Beyond TBI, several investigations have supported an association between tau pathology and seizures (Sanchez et al., 2018; Tai et al., 2016). Studies on epileptic human temporal structure revealed accumulation of tau aggregates (Sanchez et al., 2018). In 3XTg AD mice, induced chronic epilepsy was associated with changes of inter-neuronal p-tau expression (Yan et al., 2012). Additionally, data obtained from post-mortem analysis of patient tissues with AD and drug resistant epilepsy uncovered a correlation between symptomatic seizures, increased Braak staging and accelerated tau accumulation (Thom et al., 2011). Interestingly, the presence of tau deposits in epileptic patients and the similarity of its pathology to CTE suggest a conceivable role for seizures influencing the progression of tau pathology in a similar manner to TBI (Puvenna et al., 2016). Indeed, our data from the application of the convulsant kainate here support the role of post-traumatic seizure in enhancing tau abundance and cell death in our TBI model (Fig. 5G,H). This finding is in line with observations in a patient with epilepsy and a history of head injury, in which progressive tau pathology was noted (Geddes et al., 1999; Thom et al., 2011). Intriguingly, reducing seizure activity after TBI via anti-convulsant drugs was able to significantly reduce tauopathy and cell death, providing further evidence of the relationship between seizures and tauopathy in TBI (Fig 5F,G and H). The mechanism of drug action appears to be dominated by its anticonvulsant properties, because its effects were reversed by co-application of convulsants. Thus, anti-convulsants are intriguing as a route to slowing progression of tauopathy following TBI, and it is encouraging that they are already commonly deployed to prevent post-traumatic seizures.

### Endocytosis mediates prion-like spread of tauopathy

One particular convulsant drug, 4-AP, inhibited tauopathy in our TBI model (Fig 6 A and B), contrary to our hypothesis that seizure intensity is positively correlated with tauopathy following TBI. 4-AP is a voltage-gated potassium channel blocker that enhances neuronal firing activity and has been used often in zebrafish seizure studies (Kasatkina, 2016; Liu and Baraban, 2019; Lundh, 1978; R Kanyo, IN REVISION Jan 16th, 2020 ; Winter et al., 2017). Yet, 4-AP is rarely administered for prolonged treatments such as those we deployed here, e.g. past studies rarely exceed one hour of 4-AP (Winter et al., 2017). Thus, we considered that our high dose and prolonged stimulation with 4-AP may have led to off-target effects; we confirmed this insomuch that the 4-AP’s inhibition of tauopathy was not related to its convulsant properties (as determined by 4-AP’s effects being unaltered by potent anti-convulsants (Fig. 6B)). Indeed previous *in vitro* work revealed high concentrations or prolonged stimulation with 4-AP has off-target effects via inhibiting dynamin, which is important for the endocytosis of synaptic vesicles at the nerve terminals (Cousin and Robinson, 2000). The inhibition of endocytosis observed in that study was independent of 4-AP-dependent seizure activity.

We further queried the potential role of dynamin-dependent endocytosis in the prion-like progression of tau pathology after TBI by applying endocytosis inhibitors that target dynamin. Dynamin is a GTPase involved in two mechanisms of endocytosis that are important for synaptic vesicle transport (Singh et al., 2017). Empirical work on human stem cell-derived neurons has indicated that tau aggregates are internalized via dynamin-dependent endocytosis and that blocking other endocytosis pathways independent of dynamin, such as bulk endocytosis and macropinocytosis, did not disrupt tau uptake (Evans et al., 2018). On the contrary, inhibiting dynamin significantly decreased the internalization of tau aggregates. Our results are in line with the previously mentioned findings that show tau progression in TBI models depends on dynamin-dependent endocytic pathways - blocking them with two different inhibitors (and with 4-AP) dramatically lessened the abundance of tau seeds (Fig 6C-F). Hence, our findings not only provide *in vivo* validation of past *in vitro* works, but also suggest mechanisms underlying prion-like spreading of tau seeds in TBI and CTE that could aid in developing therapeutic strategies.

### Limitations of our approach

Our tauopathy biosensor, human Tau4R-GFP, was deployed *in vivo* and uniquely able to detect significant increases (and decreases) in the abundance of tau aggregates following various insults and treatments, typically in a dose-dependent manner and in harmony with expected trends. Considering this success, it remains intriguing that a subset of larvae exhibit GFP+ tau puncta despite receiving no known tauopathy-inducing insults. This suggests the larvae express the transgene at a level near to a threshold for producing spontaneous aggregates. We performed selective breeding to minimize these occurrences and tentatively believe, after too few generations, that genetics of the fish is a factor – substantial genetic variation exists in zebrafish in-bred lines (Balik-Meisner et al., 2018; Guryev et al., 2006). However, we acknowledge the variation could be a minor technical artefact rather than biological. More optimistically, this inter-individual variation and stochastic appearance of tauopathy, in a high-throughput model, could be leveraged to newly appreciate aspects of spontaneous AD or other non-familial tauopathies. Regardless, future work will also need to characterize the biochemistry and biophysics of the human Tau inclusions in zebrafish compared to patients or rodent models.

Regarding our TBI methods, further refinements may yet be able to improve consistency of the injury and reduce the apparent variability between individuals. This variability is real, but somewhat offset by the large sample sizes attainable: our TBI methods offer the potent advantages of zebrafish larvae with respect to genetic and drug accessibility in high-throughput formats, while also retaining the critical *in vivo* complexity required to investigate disease etiology and treatments. Further, it remains to be established if the mechanisms we reveal are ubiquitous across the various forms of TBI: our model fills a gap by supplying a rare ‘closed head’ TBI model (as opposed to the majority of animal models that access the brain by removing skull elements prior to brain injury, see exceptions by (Meconi et al., 2018; Mychasiuk et al., 2014). Our model might be most relevant to brain trauma experienced by the human foetus (e.g. during car collisions or domestic abuse), considering the developmental stage and aqueous media. Further work is also needed to appreciate how the physics of our blast injury is altered by occurring at a small scale (e.g. larval brain is <500 µm). At this point we are left to assume that the cellular and physiological aspects of TBI we consider here are sufficiently similar across all classes of TBI, and thus the knowledge gleaned may be variably applicable.

Finally, we have chosen to restrict our analysis to study of larval fish. While this offers many logistical and ethical advantages detailed above, it limits our study to acute effects occurring over the course of several days. Conclusions from such work, once refined and validated using the power of the *in vivo* zebrafish larva model, should be tested in rodent models where it is equally time-consuming to assess the long-term efficacy of treatments on these progressive late-onset dementias.

### Conclusion

Currently, no available treatments are applicable to all tauopathies, which remain as devastating and inevitably fatal dementias. Zebrafish larvae, fostered by appropriate innovations, now offer a potent complement both to rodent models of TBI and to cellular models of tauopathy. Our engineered fish allowed us to reveal post-traumatic seizures as a druggable mechanistic link between TBI and the prion-like progression of tauopathy. Intriguingly, our conclusions have potential for translation to TBI clinics where anti-convulsants are already in use as prophylactics for post-traumatic epilepsy, though further work remains to address if they mitigate (the risk or severity of) later progression of CTE, AD or other tauopathies.

## Acknowledgements

We acknowledge Nathalie Daude and David Westaway provided mouse brain samples, advice, and access to cell culture infrastructure. Gavin Neil and Jenna Bratvold contributed to SOD1-GFP cloning and zebrafish transgenesis via modifying a vector provided by Neil Cashman and Edward Pokrishevsky. Mark Loewen provided advice on hydraulic measures of pressure. Sue-Ann Mok, Satya Kar, Oksana Suchowersky, David Westaway and Brian Christie provided comments on an earlier version of the manuscript.

Funding to HA was from the Saudi Arabia Cultural Bureau and Majmaah University. RK was supported by SynAD postdoctoral fellowship funded via Alzheimer Society of Alberta and Northwest Territories through their Hope for Tomorrow program and the University Hospital Foundation. LFL received Studentships from Alberta Innovates and NSERC. MGD received Studentships from Alberta Innovates and CIHR. Operating funds to EB were from CurePSP 655-2018-06 and 468-08, US Department of Veterans Affairs BX003168, and NIH NS080881; The contents of this article do not represent the views of the United States government. Operating funds to HW were from Alberta Innovates and the Alzheimer Society of Alberta and Northwest Territories through the joint Alberta Alzheimer’s Research Program (AARP 201700005). Operating funds to WTA were also from the joint AARP (201700018), and from anonymous donors; donors played no role in study design, prioritization or data interpretation, or decision to publish.

## Author Contributions

HA preformed the experiments, collected and analyzed data, composed and wrote the manuscript. RK collected and analyzed data for CaMPARI experiments. LFL preformed kainate and some retigabine experiments, and characterized pressure kinetics. RK-J and HW obtained the electron microscope images. QB and EAB engineered and provided human Tau Tg zebrafish. MGD engineered SOD1:GFP Tg zebrafish. WTA was the supervisory author, was involved in concept formation, data analysis and interpretation, and edited the manuscript. All authors advised on editing the manuscript.

## References

Asikainen, I., Kaste, M., and Sarna, S. (1999). Early and late posttraumatic seizures in traumatic brain injury rehabilitation patients: brain injury factors causing late seizures and influence of seizures on long-term outcome. Epilepsia 40, 584–589.

Ayers, J.I., Giasson, B.I., and Borchelt, D.R. (2018). Prion-like Spreading in Tauopathies. Biol Psychiatry 83, 337–346.

Bai, Q., Garver, J.A., Hukriede, N.A., and Burton, E.A. (2007). Generation of a transgenic zebrafish model of Tauopathy using a novel promoter element derived from the zebrafish eno2 gene. Nucleic Acids Res 35, 6501–6516.

Balik-Meisner, M., Truong, L., Scholl, E.H., Tanguay, R.L., and Reif, D.M. (2018). Population genetic diversity in zebrafish lines. Mamm Genome 29, 90–100.

Bir, C., Vandevord, P., Shen, Y., Raza, W., and Haacke, E.M. (2012). Effects of variable blast pressures on blood flow and oxygen saturation in rat brain as evidenced using MRI. Magn Reson Imaging 30, 527–534.

Bodnar, C.N., Roberts, K.N., Higgins, E.K., and Bachstetter, A.D. (2019). A Systematic Review of Closed Head Injury Models of Mild Traumatic Brain Injury in Mice and Rats. J Neurotrauma 36, 1683–1706.

Bugay, V., Bozdemir, E., Vigil, F.A., Holstein, D.M., Chun, S.H., Elliot, W., Sprague, C., Cavazos, J.E., Zamora, D.O., Rule, G., et al. (2019). A mouse model of repetitive blast traumatic brain injury reveals post-trauma seizures and increased neuronal excitability. J Neurotrauma.

Chauhan, N.B. (2014). Chronic neurodegenerative consequences of traumatic brain injury. Restor Neurol Neurosci 32, 337–365.

Chiti, F., and Dobson, C.M. (2006). Protein misfolding, functional amyloid, and human disease. Annu Rev Biochem 75, 333–366.

Clavaguera, F., Akatsu, H., Fraser, G., Crowther, R.A., Frank, S., Hench, J., Probst, A., Winkler, D.T., Reichwald, J., Staufenbiel, M., et al. (2013). Brain homogenates from human tauopathies induce tau inclusions in mouse brain. Proc Natl Acad Sci U S A 110, 9535–9540.

Colin, M., Dujardin, S., Schraen-Maschke, S., Meno-Tetang, G., Duyckaerts, C., Courade, J.P., and Buee, L. (2020). From the prion-like propagation hypothesis to therapeutic strategies of anti-tau immunotherapy. Acta Neuropathol 139, 3–25.

Cousin, M.A., and Robinson, P.J. (2000). Ca(2+) influx inhibits dynamin and arrests synaptic vesicle endocytosis at the active zone. J Neurosci 20, 949–957.

Cruz-Haces, M., Tang, J., Acosta, G., Fernandez, J., and Shi, R. (2017). Pathological correlations between traumatic brain injury and chronic neurodegenerative diseases. Transl Neurodegener 6, 20.

de Calignon, A., Polydoro, M., Suarez-Calvet, M., William, C., Adamowicz, D.H., Kopeikina, K.J., Pitstick, R., Sahara, N., Ashe, K.H., Carlson, G.A., et al. (2012). Propagation of tau pathology in a model of early Alzheimer’s disease. Neuron 73, 685–697.

Demaegd, K., Schymkowitz, J., and Rousseau, F. (2018). Transcellular Spreading of Tau in Tauopathies. Chembiochem 19, 2424–2432.

DuVal, M.G., Oel, A.P., and Allison, W.T. (2014). gdf6a Is Required for Cone Photoreceptor Subtype Differentiation and for the Actions of tbx2b in Determining Rod Versus Cone Photoreceptor Fate. Plos One 9.

Ellis, L.D., Seibert, J., and Soanes, K.H. (2012). Distinct models of induced hyperactivity in zebrafish larvae. Brain Res 1449, 46–59.

Englander, J., Cifu, D.X., and Diaz-Arrastia, R. (2014). Information/education page. Seizures and traumatic brain injury. Arch Phys Med Rehabil 95, 1223–1224.

Eskandari-Sedighi, G., Daude, N., Gapeshina, H., Sanders, D.W., Kamali-Jamil, R., Yang, J., Shi, B., Wille, H., Ghetti, B., Diamond, M.I., et al. (2017). The CNS in inbred transgenic models of 4-repeat Tauopathy develops consistent tau seeding capacity yet focal and diverse patterns of protein deposition. Mol Neurodegener 12, 72.

Evans, L.D., Wassmer, T., Fraser, G., Smith, J., Perkinton, M., Billinton, A., and Livesey, F.J. (2018). Extracellular Monomeric and Aggregated Tau Efficiently Enter Human Neurons through Overlapping but Distinct Pathways. Cell Rep 22, 3612–3624.

Fisher, S., Grice, E.A., Vinton, R.M., Bessling, S.L., Urasaki, A., Kawakami, K., and McCallion, A.S. (2006). Evaluating the biological relevance of putative enhancers using Tol2 transposon-mediated transgenesis in zebrafish. Nat Protoc 1, 1297–1305.

Fosque, B.F., Sun, Y., Dana, H., Yang, C.T., Ohyama, T., Tadross, M.R., Patel, R., Zlatic, M., Kim, D.S., Ahrens, M.B., et al. (2015). Neural circuits. Labeling of active neural circuits in vivo with designed calcium integrators. Science 347, 755–760.

Gardner, R.C., and Yaffe, K. (2015). Epidemiology of mild traumatic brain injury and neurodegenerative disease. Mol Cell Neurosci 66, 75–80.

Geddes, J.F., Vowles, G.H., Nicoll, J.A., and Revesz, T. (1999). Neuronal cytoskeletal changes are an early consequence of repetitive head injury. Acta Neuropathol 98, 171–178.

Goedert, M., Eisenberg, D.S., and Crowther, R.A. (2017a). Propagation of Tau Aggregates and Neurodegeneration. Annu Rev Neurosci 40, 189–210.

Goedert, M., Masuda-Suzukake, M., and Falcon, B. (2017b). Like prions: the propagation of aggregated tau and alpha-synuclein in neurodegeneration. Brain 140, 266–278.

Goedert, M., Spillantini, M.G., Jakes, R., Rutherford, D., and Crowther, R.A. (1989). Multiple isoforms of human microtubule-associated protein tau: sequences and localization in neurofibrillary tangles of Alzheimer’s disease. Neuron 3, 519–526.

Guo, J.L., and Lee, V.M. (2011). Seeding of normal Tau by pathological Tau conformers drives pathogenesis of Alzheimer-like tangles. J Biol Chem 286, 15317–15331.

Guo, J.L., Narasimhan, S., Changolkar, L., He, Z., Stieber, A., Zhang, B., Gathagan, R.J., Iba, M., McBride, J.D., Trojanowski, J.Q., and Lee, V.M. (2016). Unique pathological tau conformers from Alzheimer’s brains transmit tau pathology in nontransgenic mice. J Exp Med 213, 2635–2654.

Guryev, V., Koudijs, M.J., Berezikov, E., Johnson, S.L., Plasterk, R.H., van Eeden, F.J., and Cuppen, E. (2006). Genetic variation in the zebrafish. Genome Res 16, 491–497.

Gutzman, J.H., and Sive, H. (2009). Zebrafish brain ventricle injection. J Vis Exp.

Hanwell, D., Hutchinson, S.A., Collymore, C., Bruce, A.E., Louis, R., Ghalami, A., Allison, W.T., Ekker, M., Eames, B.F., Childs, S., et al. (2016). Restrictions on the Importation of Zebrafish into Canada Associated with Spring Viremia of Carp Virus. Zebrafish 13 Suppl 1, S153–163.

Hay, J., Johnson, V.E., Smith, D.H., and Stewart, W. (2016). Chronic Traumatic Encephalopathy: The Neuropathological Legacy of Traumatic Brain Injury. Annu Rev Pathol 11, 21–45.

Iba, M., Guo, J.L., McBride, J.D., Zhang, B., Trojanowski, J.Q., and Lee, V.M. (2013). Synthetic tau fibrils mediate transmission of neurofibrillary tangles in a transgenic mouse model of Alzheimer’s-like tauopathy. J Neurosci 33, 1024–1037.

Iba, M., McBride, J.D., Guo, J.L., Zhang, B., Trojanowski, J.Q., and Lee, V.M. (2015). Tau pathology spread in PS19 tau transgenic mice following locus coeruleus (LC) injections of synthetic tau fibrils is determined by the LC’s afferent and efferent connections. Acta Neuropathol 130, 349–362.

Johnson, V.E., Stewart, J.E., Begbie, F.D., Trojanowski, J.Q., Smith, D.H., and Stewart, W. (2013). Inflammation and white matter degeneration persist for years after a single traumatic brain injury. Brain 136, 28–42.

Johnson, V.E., Stewart, W., and Smith, D.H. (2012). Widespread tau and amyloid-beta pathology many years after a single traumatic brain injury in humans. Brain Pathol 22, 142–149.

Kanyo, R., Leighton, P.L.A., Neil, G.J., Locskai, L.F., and Allison, W.T. (2020). Amyloid-beta precursor protein mutant zebrafish exhibit seizure susceptibility that depends on prion protein. Exp Neurol, 113283.

Kanyo, R., Wang, C.K., Locskai, L.F., Li, J., Allison, W.T., and H Kurata (IN REVISION as of Jan 16^th^, 2020). Functional and behavioral signatures of Kv7 activator drug subtypes. Epilepsia. ms# EPI-00953-2019.

Kasatkina, L.A. (2016). 4-capital A, Cyrillicminopyridine sequesters intracellular Ca(2+) which triggers exocytosis in excitable and non-excitable cells. Sci Rep 6, 34749.

Kaufman, S.K., Sanders, D.W., Thomas, T.L., Ruchinskas, A.J., Vaquer-Alicea, J., Sharma, A.M., Miller, T.M., and Diamond, M.I. (2016). Tau Prion Strains Dictate Patterns of Cell Pathology, Progression Rate, and Regional Vulnerability In Vivo. Neuron 92, 796–812.

Kovacs, G.G. (2017). Tauopathies. Handb Clin Neurol 145, 355–368.

Kovacs, S.K., Leonessa, F., and Ling, G.S. (2014). Blast TBI Models, Neuropathology, and Implications for Seizure Risk. Front Neurol 5, 47.

Kwan, K.M., Fujimoto, E., Grabher, C., Mangum, B.D., Hardy, M.E., Campbell, D.S., Parant, J.M., Yost, H.J., Kanki, J.P., and Chien, C.B. (2007). The Tol2kit: a multisite gateway-based construction kit for Tol2 transposon transgenesis constructs. Dev Dyn 236, 3088–3099.

Leighton, P.L.A., Kanyo, R., Neil, G.J., Pollock, N.M., and Allison, W.T. (2018). Prion gene paralogs are dispensable for early zebrafish development and have nonadditive roles in seizure susceptibility. J Biol Chem 293, 12576–12592.

Lim, J., and Yue, Z. (2015). Neuronal aggregates: formation, clearance, and spreading. Dev Cell 32, 491–501.

Liu, J., and Baraban, S.C. (2019). Network Properties Revealed during Multi-Scale Calcium Imaging of Seizure Activity in Zebrafish. eNeuro 6.

Lucke-Wold, B.P., Nguyen, L., Turner, R.C., Logsdon, A.F., Chen, Y.W., Smith, K.E., Huber, J.D., Matsumoto, R., Rosen, C.L., Tucker, E.S., and Richter, E. (2015). Traumatic brain injury and epilepsy: Underlying mechanisms leading to seizure. Seizure 33, 13–23.

Lundh, H. (1978). Effects of 4-aminopyridine on neuromuscular transmission. Brain Res 153, 307–318.

Maheras, A.L., Dix, B., Carmo, O.M.S., Young, A.E., Gill, V.N., Sun, J.L., Booker, A.R., Thomason, H.A., Ibrahim, A.E., Stanislaw, L., et al. (2018). Genetic Pathways of Neuroregeneration in a Novel Mild Traumatic Brain Injury Model in Adult Zebrafish. eNeuro 5.

Marklund, N. (2016). Rodent Models of Traumatic Brain Injury: Methods and Challenges. Methods Mol Biol 1462, 29–46.

McCluskey, A., Daniel, J.A., Hadzic, G., Chau, N., Clayton, E.L., Mariana, A., Whiting, A., Gorgani, N.N., Lloyd, J., Quan, A., et al. (2013). Building a better dynasore: the dyngo compounds potently inhibit dynamin and endocytosis. Traffic 14, 1272–1289.

McCutcheon, V., Park, E., Liu, E., Sobhebidari, P., Tavakkoli, J., Wen, X.Y., and Baker, A.J. (2017). A Novel Model of Traumatic Brain Injury in Adult Zebrafish Demonstrates Response to Injury and Treatment Comparable with Mammalian Models. J Neurotrauma 34, 1382–1393.

McGeachie, A.B., Odell, L.R., Quan, A., Daniel, J.A., Chau, N., Hill, T.A., Gorgani, N.N., Keating, D.J., Cousin, M.A., van Dam, E.M., et al. (2013). Pyrimidyn compounds: dual-action small molecule pyrimidine-based dynamin inhibitors. ACS Chem Biol 8, 1507–1518.

McKee, A.C., Stein, T.D., Kiernan, P.T., and Alvarez, V.E. (2015). The neuropathology of chronic traumatic encephalopathy. Brain Pathol 25, 350–364.

Meconi, A., Wortman, R.C., Wright, D.K., Neale, K.J., Clarkson, M., Shultz, S.R., and Christie, B.R. (2018). Repeated mild traumatic brain injury can cause acute neurologic impairment without overt structural damage in juvenile rats. PLoS One 13, e0197187.

Mudher, A., Colin, M., Dujardin, S., Medina, M., Dewachter, I., Alavi Naini, S.M., Mandelkow, E.M., Mandelkow, E., Buee, L., Goedert, M., and Brion, J.P. (2017). What is the evidence that tau pathology spreads through prion-like propagation? Acta Neuropathol Commun 5, 99.

Murakami, T., Paitel, E., Kawarabayashi, T., Ikeda, M., Chishti, M.A., Janus, C., Matsubara, E., Sasaki, A., Kawarai, T., Phinney, A.L., et al. (2006). Cortical neuronal and glial pathology in TgTauP301L transgenic mice: neuronal degeneration, memory disturbance, and phenotypic variation. Am J Pathol 169, 1365–1375.

Mychasiuk, R., Farran, A., Angoa-Perez, M., Briggs, D., Kuhn, D., and Esser, M.J. (2014). A novel model of mild traumatic brain injury for juvenile rats. J Vis Exp.

Nakagawa, A., Manley, G.T., Gean, A.D., Ohtani, K., Armonda, R., Tsukamoto, A., Yamamoto, H., Takayama, K., and Tominaga, T. (2011). Mechanisms of primary blast-induced traumatic brain injury: insights from shock-wave research. J Neurotrauma 28, 1101–1119.

Narasimhan, S., Guo, J.L., Changolkar, L., Stieber, A., McBride, J.D., Silva, L.V., He, Z., Zhang, B., Gathagan, R.J., Trojanowski, J.Q., and Lee, V.M.Y. (2017). Pathological Tau Strains from Human Brains Recapitulate the Diversity of Tauopathies in Nontransgenic Mouse Brain. J Neurosci 37, 11406–11423.

Nguyen, R., Fiest, K.M., McChesney, J., Kwon, C.S., Jette, N., Frolkis, A.D., Atta, C., Mah, S., Dhaliwal, H., Reid, A., et al. (2016). The International Incidence of Traumatic Brain Injury: A Systematic Review and Meta-Analysis. Can J Neurol Sci 43, 774–785.

Ojo, J.O., Mouzon, B., Algamal, M., Leary, P., Lynch, C., Abdullah, L., Evans, J., Mullan, M., Bachmeier, C., Stewart, W., and Crawford, F. (2016). Chronic Repetitive Mild Traumatic Brain Injury Results in Reduced Cerebral Blood Flow, Axonal Injury, Gliosis, and Increased T-Tau and Tau Oligomers. J Neuropathol Exp Neurol 75, 636–655.

Orr, M.E., Sullivan, A.C., and Frost, B. (2017). A Brief Overview of Tauopathy: Causes, Consequences, and Therapeutic Strategies. Trends Pharmacol Sci 38, 637–648.

Peeraer, E., Bottelbergs, A., Van Kolen, K., Stancu, I.C., Vasconcelos, B., Mahieu, M., Duytschaever, H., Ver Donck, L., Torremans, A., Sluydts, E., et al. (2015). Intracerebral injection of preformed synthetic tau fibrils initiates widespread tauopathy and neuronal loss in the brains of tau transgenic mice. Neurobiol Dis 73, 83–95.

Pham, L., Shultz, S.R., Kim, H.A., Brady, R.D., Wortman, R.C., Genders, S.G., Hale, M.W., O’Shea, R.D., Djouma, E., van den Buuse, M., et al. (2019). Mild Closed-Head Injury in Conscious Rats Causes Transient Neurobehavioral and Glial Disturbances: A Novel Experimental Model of Concussion. J Neurotrauma 36, 2260–2271.

Pickett, E.K., Henstridge, C.M., Allison, E., Pitstick, R., Pooler, A., Wegmann, S., Carlson, G., Hyman, B.T., and Spires-Jones, T.L. (2017). Spread of tau down neural circuits precedes synapse and neuronal loss in the rTgTauEC mouse model of early Alzheimer’s disease. Synapse 71.

Ping, X., and Jin, X. (2016). Transition from Initial Hypoactivity to Hyperactivity in Cortical Layer V Pyramidal Neurons after Traumatic Brain Injury In Vivo. J Neurotrauma 33, 354–361.

Pokrishevsky, E., McAlary, L., Farrawell, N.E., Zhao, B., Sher, M., Yerbury, J.J., and Cashman, N.R. (2018). Tryptophan 32-mediated SOD1 aggregation is attenuated by pyrimidine-like compounds in living cells. Sci Rep 8, 15590.

Puvenna, V., Engeler, M., Banjara, M., Brennan, C., Schreiber, P., Dadas, A., Bahrami, A., Solanki, J., Bandyopadhyay, A., Morris, J.K., et al. (2016). Is phosphorylated tau unique to chronic traumatic encephalopathy? Phosphorylated tau in epileptic brain and chronic traumatic encephalopathy. Brain Res 1630, 225–240.

Rimel, R.W., Giordani, B., Barth, J.T., Boll, T.J., and Jane, J.A. (1981). Disability caused by minor head injury. Neurosurgery 9, 221–228.

Russell, W.M.S., and Burch, R.L. (1959). The principles of humane experimental technique (London,: Methuen).

Safar, J., Wille, H., Itri, V., Groth, D., Serban, H., Torchia, M., Cohen, F.E., and Prusiner, S.B. (1998). Eight prion strains have PrP(Sc) molecules with different conformations. Nat Med 4, 1157–1165.

Saleem, S., and Kannan, R.R. (2018). Zebrafish: an emerging real-time model system to study Alzheimer’s disease and neurospecific drug discovery. Cell Death Discov 4, 45.

Salinsky, M., Storzbach, D., Goy, E., and Evrard, C. (2015). Traumatic brain injury and psychogenic seizures in veterans. J Head Trauma Rehabil 30, E65–70.

Sanchez, M.P., Garcia-Cabrero, A.M., Sanchez-Elexpuru, G., Burgos, D.F., and Serratosa, J.M. (2018). Tau-Induced Pathology in Epilepsy and Dementia: Notions from Patients and Animal Models. Int J Mol Sci 19.

Sanders, D.W., Kaufman, S.K., DeVos, S.L., Sharma, A.M., Mirbaha, H., Li, A., Barker, S.J., Foley, A.C., Thorpe, J.R., Serpell, L.C., et al. (2014). Distinct tau prion strains propagate in cells and mice and define different tauopathies. Neuron 82, 1271–1288.

Singh, M., Jadhav, H.R., and Bhatt, T. (2017). Dynamin Functions and Ligands: Classical Mechanisms Behind. Mol Pharmacol 91, 123–134.

Stafstrom, C.E., and Carmant, L. (2015). Seizures and epilepsy: an overview for neuroscientists. Cold Spring Harb Perspect Med 5.

Tai, X.Y., Koepp, M., Duncan, J.S., Fox, N., Thompson, P., Baxendale, S., Liu, J.Y., Reeves, C., Michalak, Z., and Thom, M. (2016). Hyperphosphorylated tau in patients with refractory epilepsy correlates with cognitive decline: a study of temporal lobe resections. Brain 139, 2441–2455.

Thom, M., Liu, J.Y., Thompson, P., Phadke, R., Narkiewicz, M., Martinian, L., Marsdon, D., Koepp, M., Caboclo, L., Catarino, C.B., and Sisodiya, S.M. (2011). Neurofibrillary tangle pathology and Braak staging in chronic epilepsy in relation to traumatic brain injury and hippocampal sclerosis: a post-mortem study. Brain 134, 2969–2981.

Traver, D., Paw, B.H., Poss, K.D., Penberthy, W.T., Lin, S., and Zon, L.I. (2003). Transplantation and in vivo imaging of multilineage engraftment in zebrafish bloodless mutants. Nat Immunol 4, 1238–1246.

Uryu, K., Chen, X.H., Martinez, D., Browne, K.D., Johnson, V.E., Graham, D.I., Lee, V.M., Trojanowski, J.Q., and Smith, D.H. (2007). Multiple proteins implicated in neurodegenerative diseases accumulate in axons after brain trauma in humans. Exp Neurol 208, 185–192.

Verweij, F.J., Revenu, C., Arras, G., Dingli, F., Loew, D., Pegtel, D.M., Follain, G., Allio, G., Goetz, J.G., Zimmermann, P., et al. (2019). Live Tracking of Inter-organ Communication by Endogenous Exosomes In Vivo. Dev Cell 48, 573–589 e574.

Wang, Y., Balaji, V., Kaniyappan, S., Kruger, L., Irsen, S., Tepper, K., Chandupatla, R., Maetzler, W., Schneider, A., Mandelkow, E., and Mandelkow, E.M. (2017). The release and trans-synaptic transmission of Tau via exosomes. Mol Neurodegener 12, 5.

Washington, P.M., Villapol, S., and Burns, M.P. (2016). Polypathology and dementia after brain trauma: Does brain injury trigger distinct neurodegenerative diseases, or should they be classified together as traumatic encephalopathy? Exp Neurol 275 Pt 3, 381–388.

Westerfield, M. (2000). The zebrafish book. A guide for the laboratory use of zebrafish (Danio rerio), 4th edn (University of Oregon press, Eugene).

White, R.M., Sessa, A., Burke, C., Bowman, T., LeBlanc, J., Ceol, C., Bourque, C., Dovey, M., Goessling, W., Burns, C.E., and Zon, L.I. (2008). Transparent adult zebrafish as a tool for in vivo transplantation analysis. Cell Stem Cell 2, 183–189.

Winter, M.J., Windell, D., Metz, J., Matthews, P., Pinion, J., Brown, J.T., Hetheridge, M.J., Ball, J.S., Owen, S.F., Redfern, W.S., et al. (2017). 4-dimensional functional profiling in the convulsant-treated larval zebrafish brain. Sci Rep 7, 6581.

Woerman, A.L., Aoyagi, A., Patel, S., Kazmi, S.A., Lobach, I., Grinberg, L.T., McKee, A.C., Seeley, W.W., Olson, S.H., and Prusiner, S.B. (2016). Tau prions from Alzheimer’s disease and chronic traumatic encephalopathy patients propagate in cultured cells. Proc Natl Acad Sci U S A 113, E8187–E8196.

Wu, J.W., Herman, M., Liu, L., Simoes, S., Acker, C.M., Figueroa, H., Steinberg, J.I., Margittai, M., Kayed, R., Zurzolo, C., et al. (2013). Small misfolded Tau species are internalized via bulk endocytosis and anterogradely and retrogradely transported in neurons. J Biol Chem 288, 1856–1870.

Wu, J.W., Hussaini, S.A., Bastille, I.M., Rodriguez, G.A., Mrejeru, A., Rilett, K., Sanders, D.W., Cook, C., Fu, H., Boonen, R.A., et al. (2016). Neuronal activity enhances tau propagation and tau pathology in vivo. Nat Neurosci 19, 1085–1092.

Yamada, K., Holth, J.K., Liao, F., Stewart, F.R., Mahan, T.E., Jiang, H., Cirrito, J.R., Patel, T.K., Hochgrafe, K., Mandelkow, E.M., and Holtzman, D.M. (2014). Neuronal activity regulates extracellular tau in vivo. J Exp Med 211, 387–393.

Yan, X.X., Cai, Y., Shelton, J., Deng, S.H., Luo, X.G., Oddo, S., Laferla, F.M., Cai, H., Rose, G.M., and Patrylo, P.R. (2012). Chronic temporal lobe epilepsy is associated with enhanced Alzheimer-like neuropathology in 3xTg-AD mice. Plos One 7, e48782.

Zanier, E.R., Bertani, I., Sammali, E., Pischiutta, F., Chiaravalloti, M.A., Vegliante, G., Masone, A., Corbelli, A., Smith, D.H., Menon, D.K., et al. (2018). Induction of a transmissible tau pathology by traumatic brain injury. Brain 141, 2685–2699.

