## Supplemental Figures for "Seizures are a druggable mechanistic link between TBI and subsequent tauopathy"

Consisting of 10 Supplemental Figures.

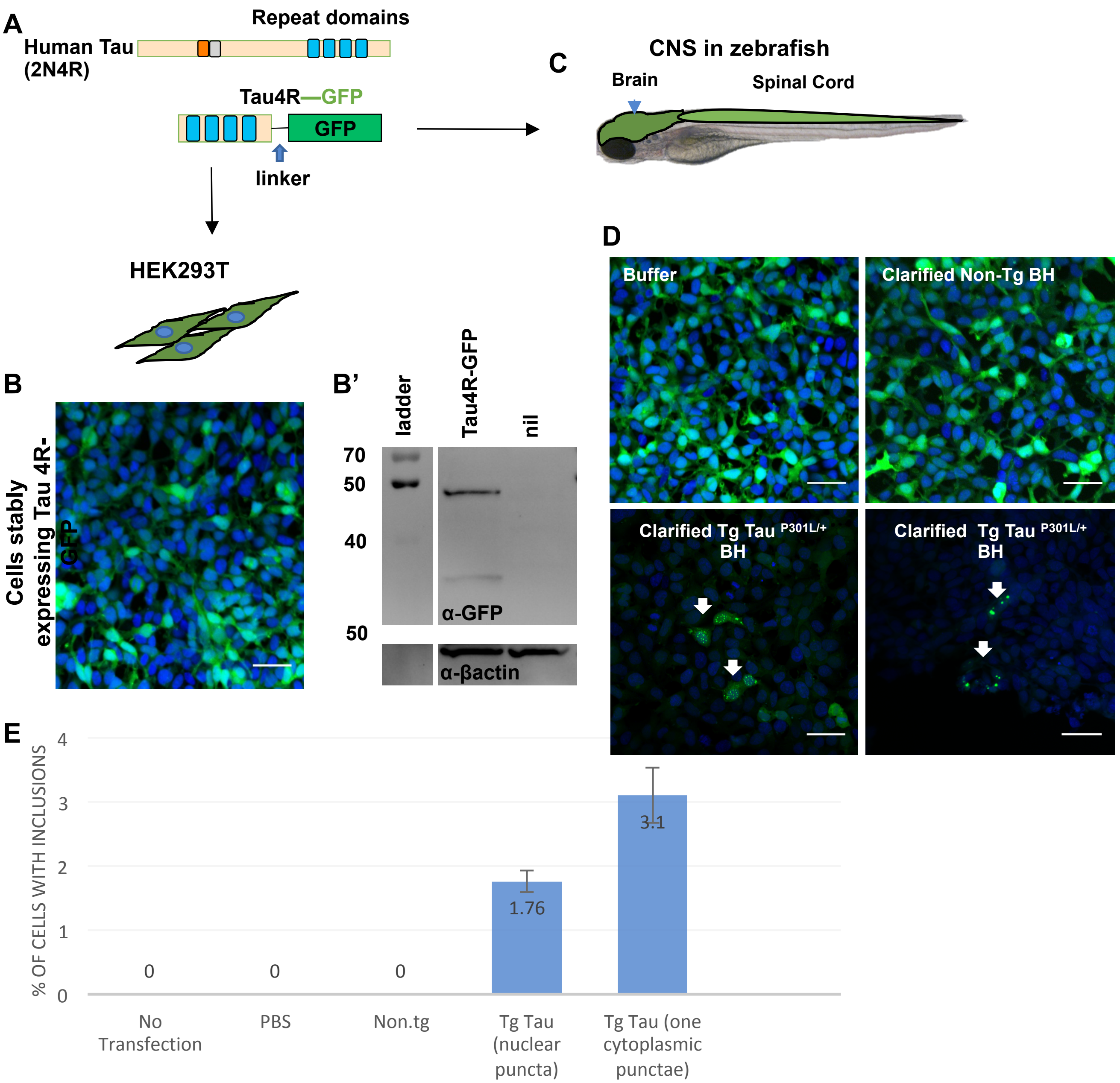


**Figure S1: Quantification GFP+ inclusions in HEK cells expressing Tau4R-GFP biosensor.** Accompanies Figure 1. **(A)** Schematic of genetically encoded fluorescent Tau4R-GFP **"Tau Biosensor"** which was produced by fusing the C-terminal four binding repeats (4R) region of wildtype human tau (Tau4R) to GFP via a linker. This construct was expressed as a transgene **(C)** in the zebrafish CNS under the promoter enolase 2, and **(B)** in Human Embryonic Kidney HEK293T cells for validation. **(B')** Immunoblot vs. GFP suggests the fusion protein Tau4R-GFP remains intact as a fusion protein when expressed in HEK cells and is an appropriate size (similar to when expressed in zebrafish CNS, Fig 1C). **(D)** Tau4R-GFP biosensor cells display GFP+ inclusions only upon transduction of crude brain homogenate burdened with tauopathy (from Tg mice expressing human Tau$^{P301L}$), but not when transduced with brain homogenate from normal mice (non-Tg BH). **(E)** HEK cells from D had a detection rate of approximately 2-3% of cells having GFP+ puncta, whereas various negative controls (no transfection, transfection with PBS only, or transfection of brain homogenate from non-Tg control mice) consistently showed no puncta. For the quantification, the number of cells with inclusions and the number of total cells from 9 field images were counted. Bars represents mean of three independent experiments.

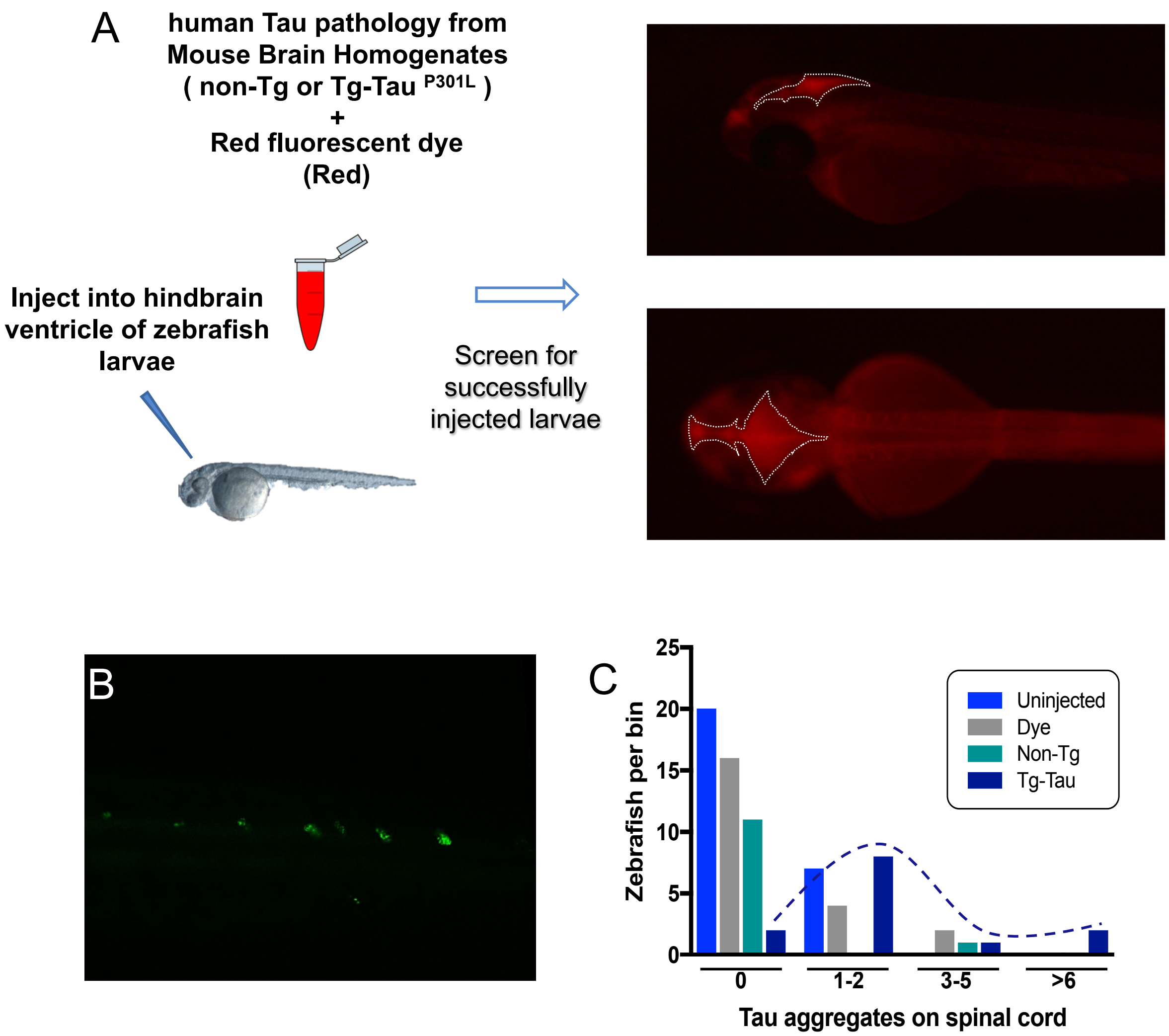


**Figure S2: Quantification GFP+ inclusions in CNS of zebrafish expressing Tau4R-GFP biosensor, following injection of human Tau into the zebrafish hindbrain ventricle.** Relates to Figure 1. **(A)** Schematic showing the microinjections of mouse brain homogenate into the brain ventricles of zebrafish larvae (age is 2 days post-fertilization). Injected material was brain homogenate from transgenic mice expressing human Tau, i.e. with tau pathology (e.g. see pathology characterized in Eskandari-Sedighi *et al.* 2017, DOI 10.1186/s13024-017-0215-7) or brain homogenate from normal control mice. Red fluorescent dye included in the diluted brain homogenate allowed imaging and selection of properly injected embryos. right: Successfully injected embryos had the injected brain homogenate localized within the zebrafish brain ventricles, with sharp edges and non-diffuse dye. The area encompassing the injected dye is outlined by white dotted lines. **(B) Similar to Tau4R-GFP aggregating in the brain, GFP+ aggregates were also detected in the spinal cord** following delivery of human Tau, akin to the brain aggregates in Figure 1F. **(C) Quantification of spinal cord aggregates.** Same data as Fig 1G re-plotted to show the distribution of larvae displaying various quantities of aggregates in the spinal cord. Colour coding of treatments equivalent to Figure 1G. Dotted line is hand drawn to emphasize the distribution of larvae injected with tau-burdened brains, compared to various the controls with peak distributions of GFP+ puncta near to zero.

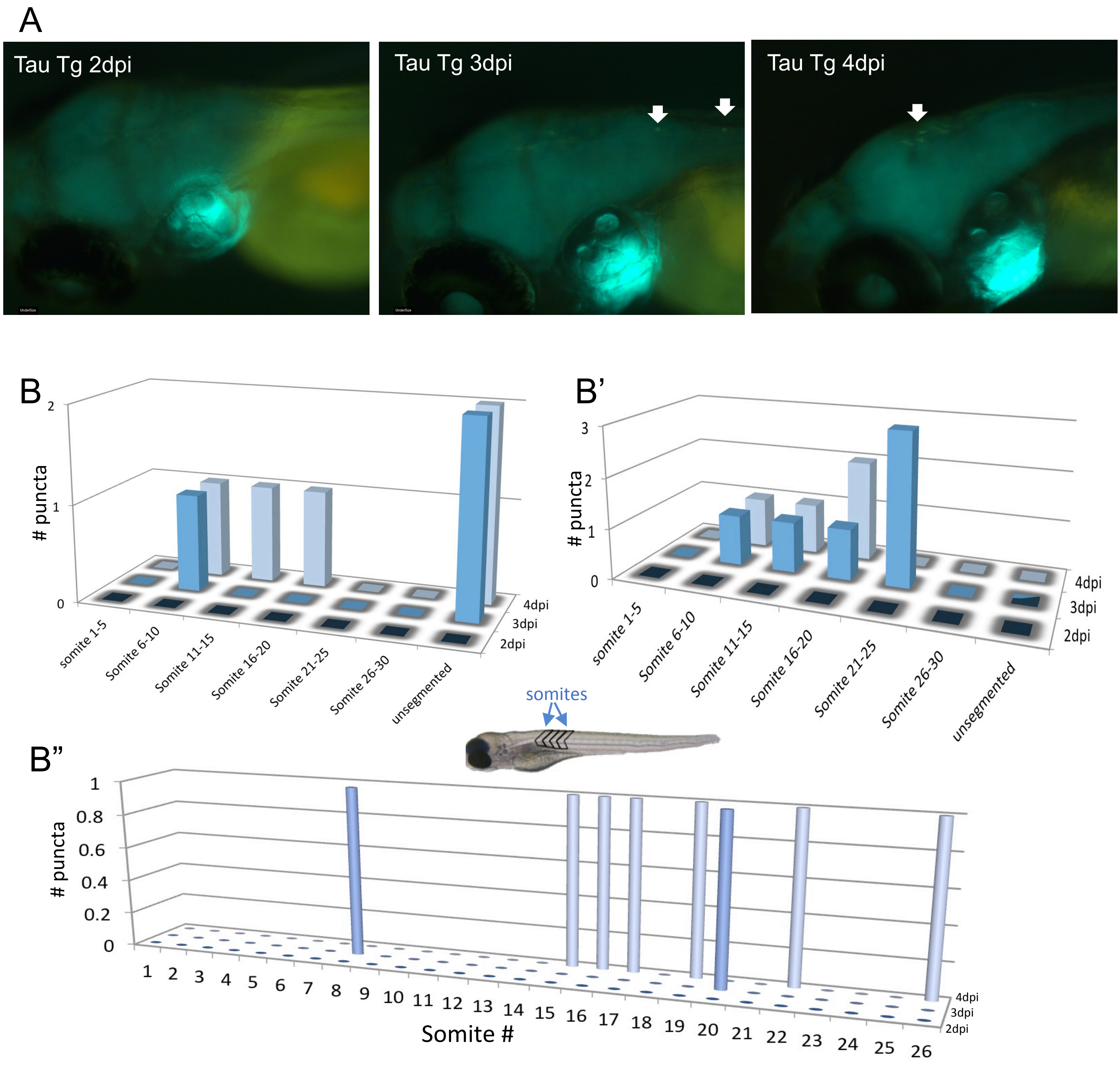


**Figure S3. Movement of some tau puncta over time following injection of zebrafish larvae with brain homogenate burdened with human tauopathy.** Relates to Figure 1. **(A)** Images of the zebrafish brain area, after injection with Tg hTau$^{+/p301L}$ mouse brain homogenate, for the same zebrafish larvae over three consecutive days post-injection (dpi), showing the movement of one puncta over time. **(B, B' and B")** The location of tau aggregates on the spinal cord of three exemplar individual larvae over multiple days, denote movement of some of these puncta over time. Somite numbers were used as landmarks to report the location along the spinal cord over multiple days - somites are blocks of tissue (e.g. muscle) that are readily visible in the trunk of the larva; larger somite number are more posterior (i.e. more distal to the injected brain region) with the most distal region being 'unsegmented'.

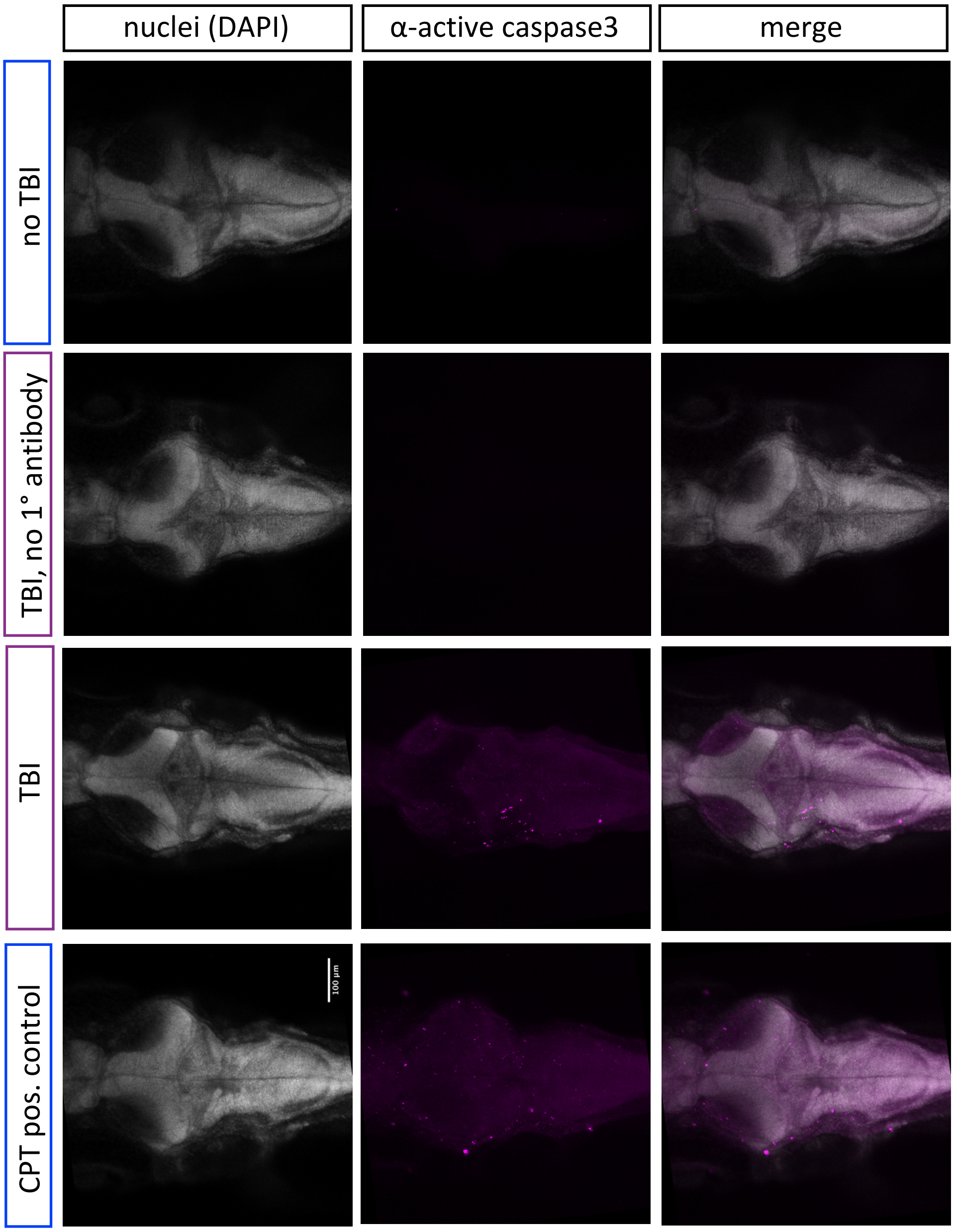


**Figure S4. Traumatic Brain Injury- (TBI-) induced cell death.** Accompanies Figure 3; some data copied from Fig 3F here for ease of reference. Increased cell death in the brain of 4 dpf larvae subjected to TBI as indicated by immunostaining of activated Caspase-3 (magenta). Larvae exposed to topoisomerase inhibitor camptothecin (CPT, 3µM) which induce apoptosis, serves as a positive control. Nuclei were stained with DAPI in gray for reference these are dorsal views of zebrafish brains with anterior at the left.

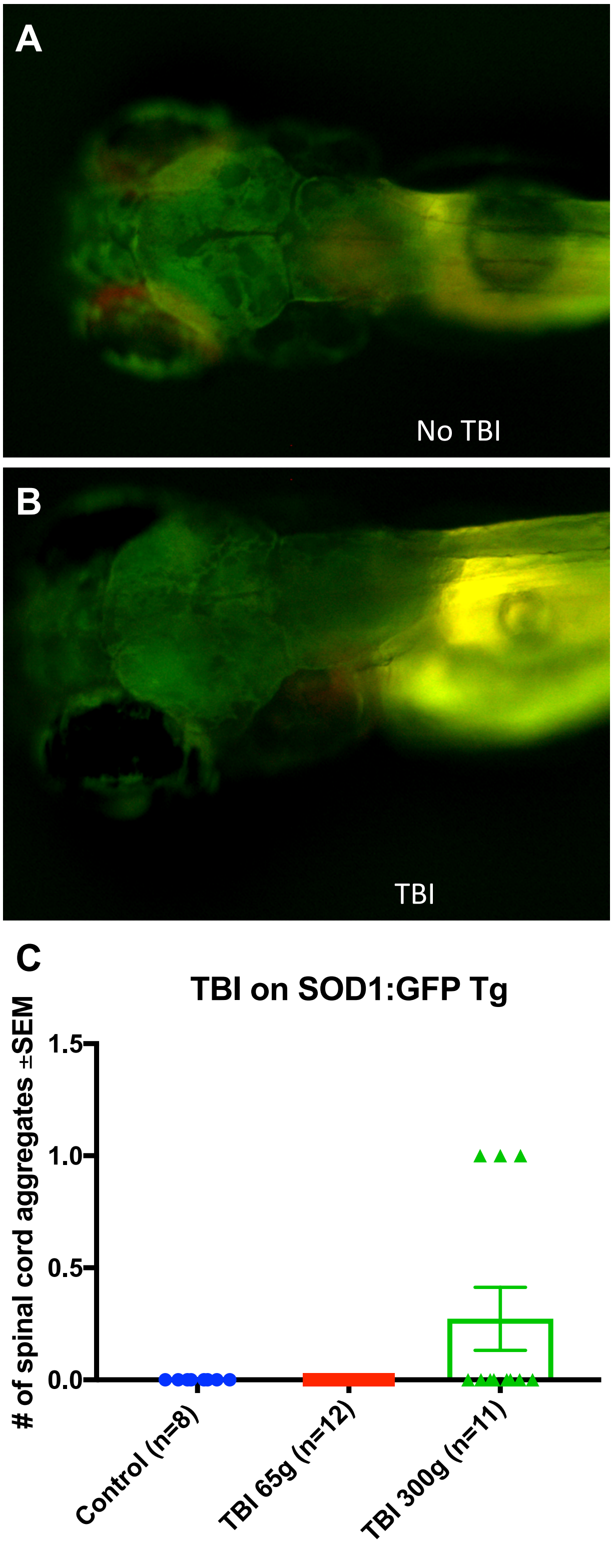


**Figure S5. Traumatic brain injury (TBI) did not induce GFP+ puncta in transgenic zebrafish larvae expressing SOD1-GFP.** Accompanies Figure 5. **(A, B)** TBI did not induce GFP+ puncta in larvae expressing SOD1-GFP, and appeared similar to control larvae that did not experience TBI. **(C)** Quantification of GFP+ puncta in the spinal cord of SOD1-GFP showing the majority of samples did not develop aggregates post traumatic brain injury.

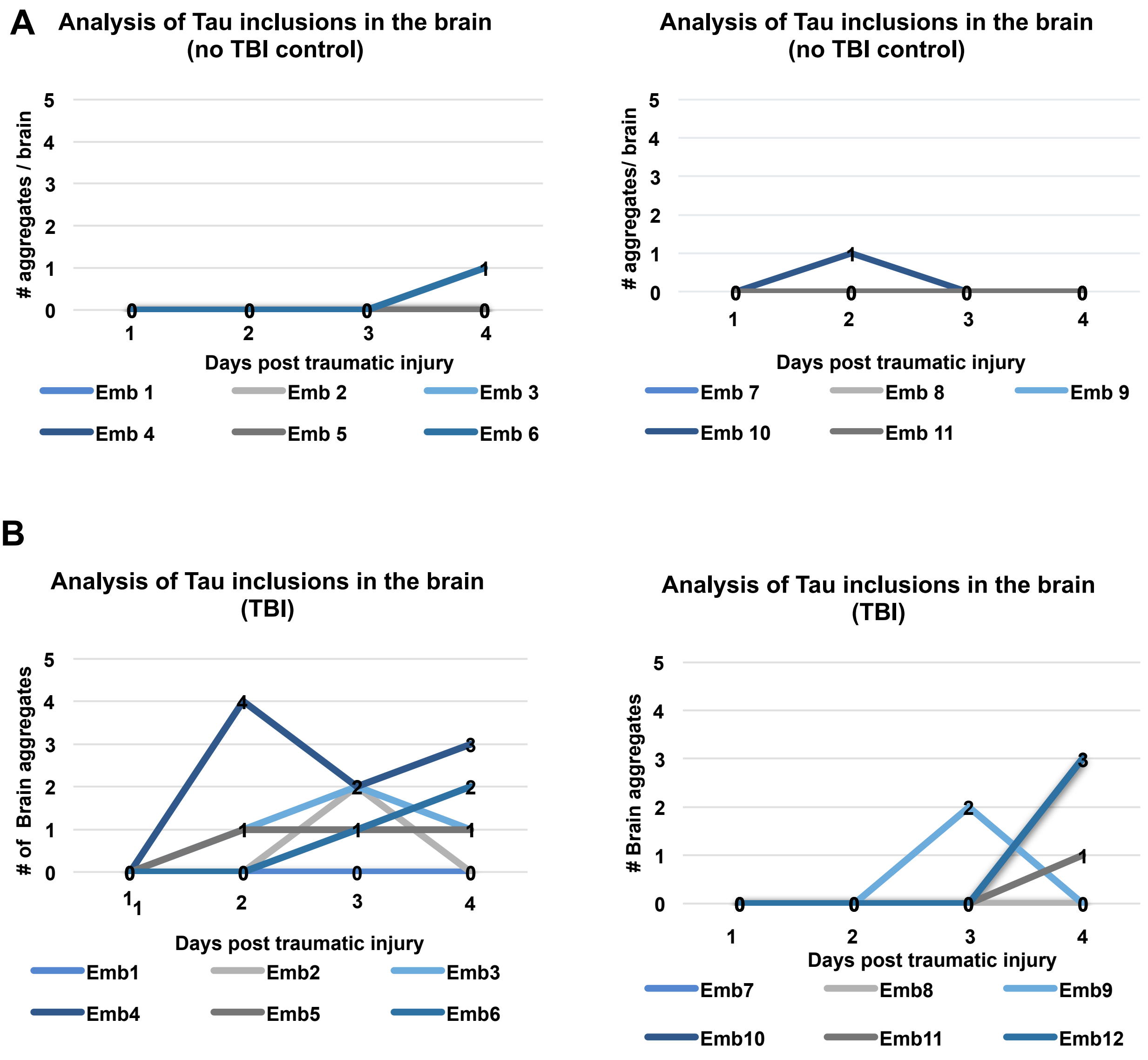


**Figure S6. Longitudinal analysis of individual fish after traumatic brain injury (TBI) shows various patterns of tau inclusion formation and clearance in their brains.** Accompanies Figure 5. **(A)** In the control group (no TBI) the majority of Tau4R-GFP biosensor larvae did not develop GFP+ aggregates in the brain, although a few either developed aggregates at later time point (3dpi), or developed aggregates that disappeared at later timepoints. **(B)** Tau4R-GFP biosensor larvae subjected to TBI developed more GFP+ brain aggregates and at earlier timepoints compared to controls, and the number of aggregates typically (though not always) increased with time. Within panels A and B, the left and right graphs are equivalent and display several larvae each.

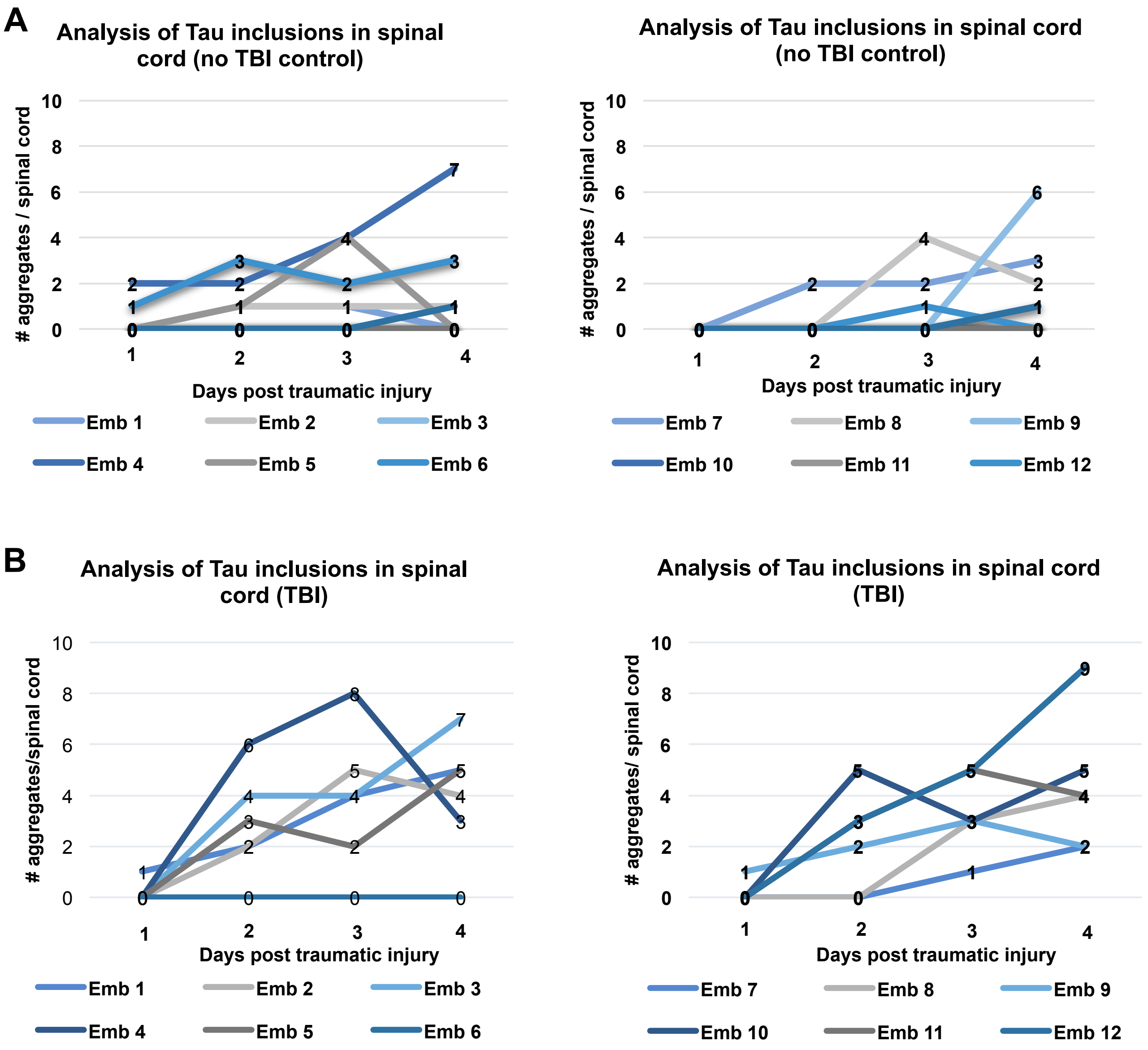


**Figure S7. Longitudinal analysis of individual fish after traumatic brain injury (TBI) shows various patterns of tau inclusion formation and clearance in their spinal cords.** Accompanies Figure 5. **(A)** Exemplar data from the control group (no TBI) wherein many of the Tau4R-GFP biosensor larvae did not develop aggregates in the spinal cord, while a few either developed aggregates at later timepoints, or developed aggregates that disappeared later on. **(B)** Tau4R-GFP biosensor larvae subjected to TBI developed more GFP+ spinal cord aggregates and at earlier timepoints compared to controls, and the number of aggregates typically (though not always) increased with time. Within panels A and B, the left and right graphs are equivalent and display several larvae each.

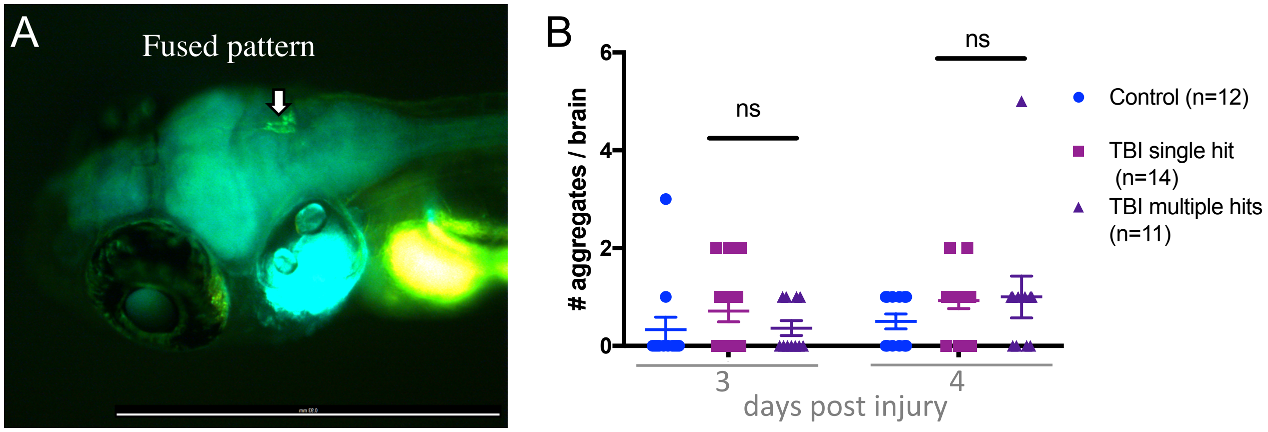


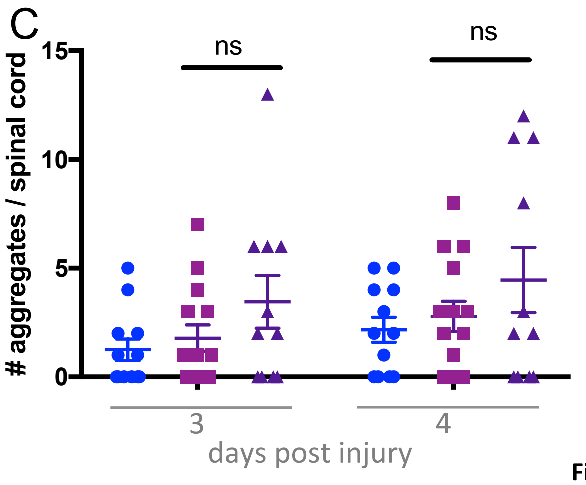


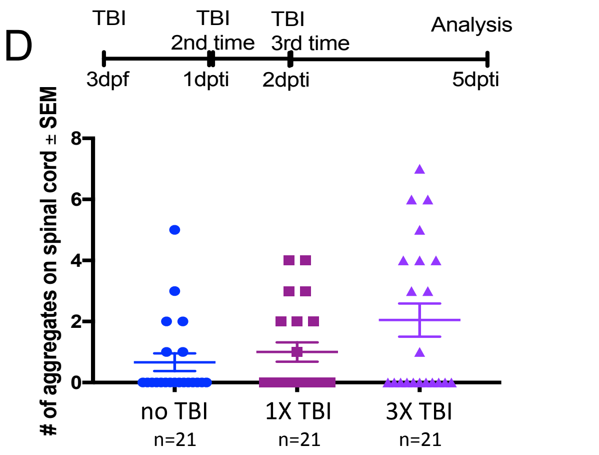


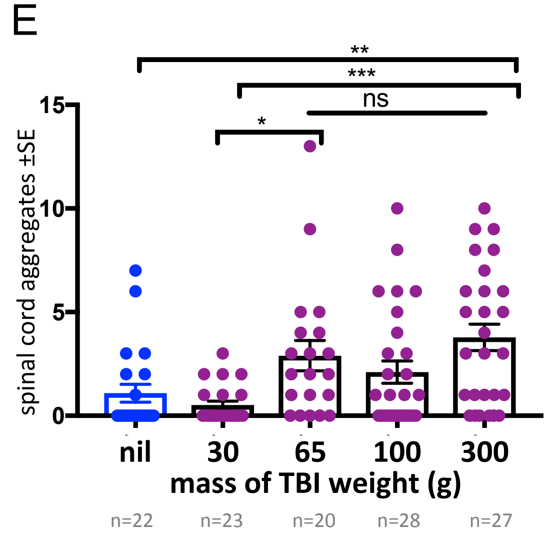


**Figure S8. Increasing intensity of traumatic brain injury (TBI) significantly increased Tau4R-GFP puncta abundance, but only modest insignificant increases in tau puncta were observed with an increasing number of successive brain injuries.** Relates to Figure 5. **(A)** Following TBI of Tau4R-GFP larvae, the GFP+ aggregates in the brain tended to be fused. **(B,C)** Tau4R-GFP larvae were subjected to TBI via either one single hit or five consecutive hits. The number of GFP+ aggregates in the brain (B) and the spinal cord (C) were quantified at 3 and 4 days post traumatic brain injury (dpi). The number of aggregates modestly increases with more hits but not consistently (not statistically significant). The symbols for treatment types are equivalent in panels B and C. **(D)** Performing TBI on Tau4R-GFP larvae on each of three consecutive days slightly increased the number of GFP+ tau puncta in the spinal cord (relative to a single TBI treatment) but the increase was not statistically significant. **(E)** A nearly linear dose-response of TBI injury intensity varying with the abundance of GFP+ puncta in the spinal cord. TBI intensity was modulated by dropping weights of varying masses in the TBI method. Heavier weights induced more GFP+ puncta (*p<0.01 and **p<0.001). In each panel raw data is plotted, along with mean ± standard error.

A

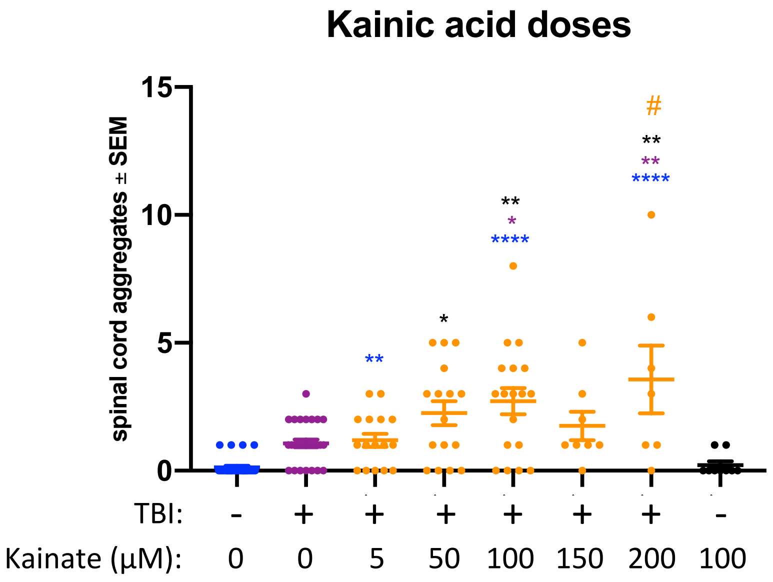


**Figure S9. Intensifying seizures following traumatic brain injury (TBI) increased abundance of GFP+ Tau puncta in a dose-dependent manner.** Relates to Figure 5. Tau4R-GFP larvae were subjected to TBI and post-traumatic seizures were intensified by addition of the convulsant kainate. Whereas kainate treatment alone (without TBI) did not appreciably increase GFP+ tau puncta in the spinal cord (black), kainate significantly increased the abundance of puncta in larvae receiving TBI (*p<0.5, **p<0.01, ****p<0.0001 compared to TBI alone [blue text]; #p<0.01 compared to TBI+5µM dose). Raw data is plotted for each larva, along with mean ± standard error.

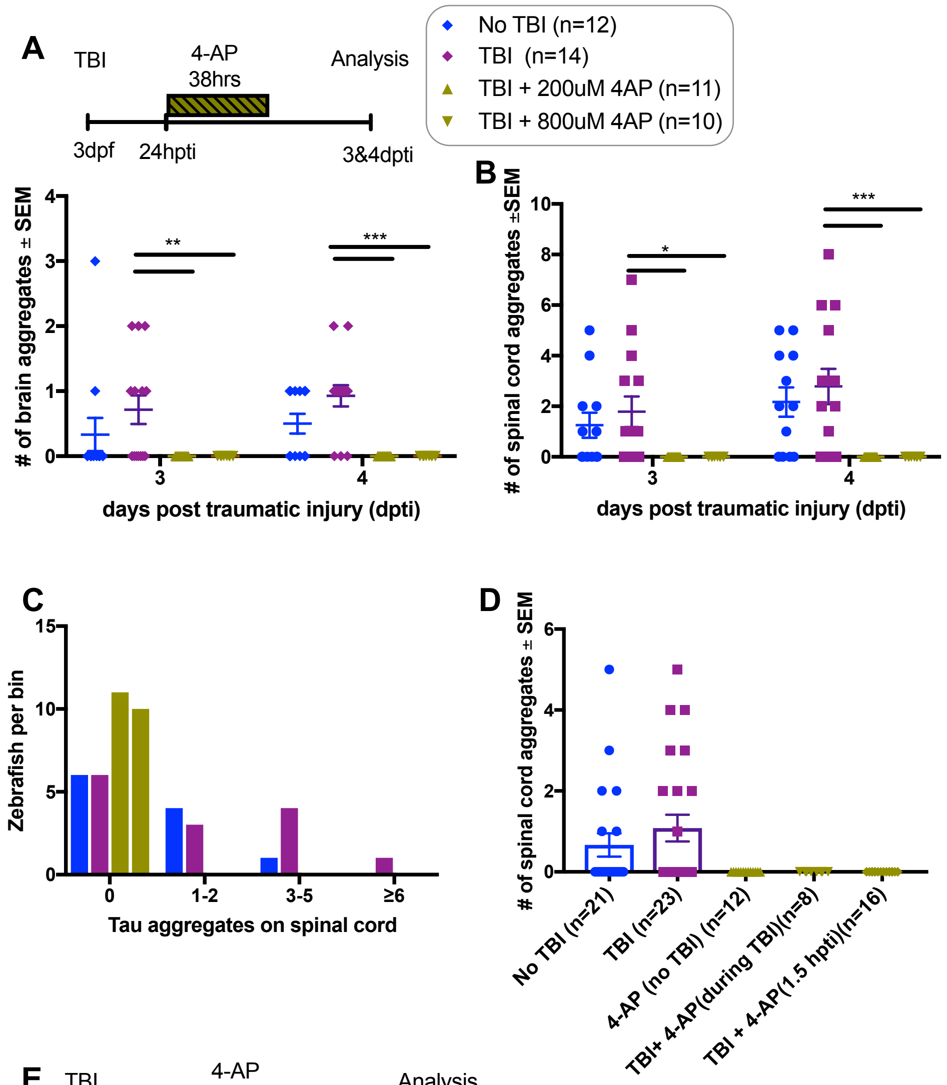


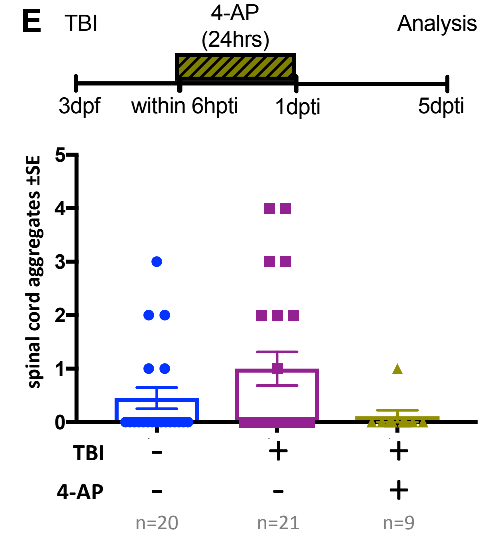


**Figure S10. 4-Aminopuridine (4-AP) abrogates TBI-induced Tau aggregates when applied at various doses or for various times.** Accompanies Figure 6. **(A)** Tau4R-GFP biosensor zebrafish larvae subjected to TBI and treated with the convulsant 4-AP show no brain puncta. 4-AP was added to the TBI larvae one day post traumatic brain injury (dpti) and left for 38 hours as displayed in timeline at top. **(A,B)** 4-AP significantly reduced (apparently eliminated) the abundance of GFP+ puncta in the brain and spinal cord compared to untreated TBI control (**$p<0.009$ at 3dpti and ***$p<0.0003$ at 4dpti) at two different doses. **(C)** All larvae treated with 4-AP failed to display any GFP+ puncta (# of aggregates is 0), displayed as distribution of larvae binned into the number of GFP+ puncta they exhibited. **(D,E)** Timing and duration of 4-AP treatments following TBI had no appreciable effect on 4-AP abrogating the TBI-induced Tau aggregates. Symbols and colours for all panels are consistent with the legend at the top right of the Figure.
